# Protein phosphatase 4 is required for centrosome asymmetry in fly neural stem cells

**DOI:** 10.1101/2025.01.15.633270

**Authors:** Roberto Carlos Segura, Emmanuel Gallaud, Adam von Barnau Sythoff, Kumar Aavula, Jennifer A. Taylor, Danielle Vahdat, Jan Pielage, Clemens Cabernard

## Abstract

Asymmetric cell division is used by stem cells to create diverse cell types while self-renewing the stem cell population. Biased segregation of molecularly distinct centrosomes could provide a mechanism to maintain stem cell fate, induce cell differentiation or both. However, the molecular mechanisms generating molecular and functional asymmetric centrosomes remain incompletely understood. Here, we show that in asymmetrically dividing fly neural stem cells, Protein phosphatase 4 (Pp4) is necessary for correct centrosome asymmetry establishment during mitosis, and microtubule organizing center (MTOC) maintenance in interphase. Using *in-vivo* live cell imaging we show that while wild type neural stem cells always maintain one active MTOC, *Pp4* mutant neuroblasts contain two inactive centrioles in interphase. Furthermore, centrosomes of *Pp4* mutant neural stem cells mature in mitosis but fail to correctly transfer the centriolar protein Centrobin (Cnb) from the mother to the daughter centriole. Using superresolution imaging, we find that phosphomimetic Centrobin fails to accurately relocalize in mitosis. We propose that Pp4 regulates the timely relocalization of Cnb in mitosis to establish two molecularly distinct centrosomes. In addition, Pp4 is also necessary to maintain MTOC activity in interphase, ensuring biased centrosome segregation. Mechanistically, Pp4 could regulate centrosome asymmetry by dephosphorylating both Cnb and gamma-Tubulin.

**SIGNIFICANCE STATEMENT:**

- Asymmetric centrosome segregation occurs in stem cells and has been linked with cell fate decisions.
- Protein phosphatase 4 (Pp4), a conserved Serine/Threonine phosphatase, regulates centrosome asymmetry in *Drosophila* neural stem cells by acting upon gamma tubulin and Centrobin.
- Pp4 regulates centrosome asymmetry establishment in mitosis and interphase, necessary for biased centrosome segregation.

## INTRODUCTION

Asymmetric cell division (ACD) is an evolutionarily conserved process that produces two cells with different fates. Stem cells routinely employ ACD to produce differentiating cells while maintaining a pool of stem cells at the same time. ACD can induce binary cell fate decisions through biased segregation of proteins, RNAs or organelles such as centrosomes (Rebollo *et al*., 2007; Yamashita *et al*., 2007; Lerit *et al*., 2013; Collins *et al*., 2017; Shlyakhtina *et al*., 2019; Delgado and Cabernard, 2020; Sunchu and Cabernard, 2020a; Gonzalez, 2021).

Centrosomes are the microtubule organization centers (MTOCs) of the cell and are composed of a pair of centrioles surrounded by a layer of pericentriolar matrix proteins (PCM). Centrioles duplicate once every cell cycle, whereby a single centriole provides a template for the formation of a younger daughter centriole. Based on this cycle, centrioles are intrinsically asymmetric by age (Conduit *et al*., 2015; Blanco-Ameijeiras *et al*., 2022). Mother and Daughter centrioles will later separate and form two new centrosomes (Conduit *et al*., 2015). Centrioles are also molecularly distinct. For instance, proteins such as Ninein, Cep194, and Outer Dense Fiber Protein 2 (ODF2) localize to the mother centriole, while Centrobin only localizes to the daughter centriole (Chen and Yamashita, 2021; Jaiswal and Singh, 2021).

Asymmetric or biased centrosome segregation has been observed in different organisms and various stem cell lineages, and the biased inheritance of centrosomes may provide a mechanistic explanation for the delivery of determinants important for cell fate decisions and development (Lambert and Nagy, 2002; Yamashita *et al*., 2007; Collins *et al*., 2017; Chen and Yamashita, 2021; Royall and Jessberger, 2021). For instance, in the developing mouse cortex, the self-renewing neuron progenitor cell (NPC) inherits the older mother centrosome, while the differentiating neuron receives the younger daughter centrosome (Wang *et al*., 2009). This is further observed in human embryonic stem cell-derived forebrain organoids, where the mother centrosome is preferentially inherited by the renewing NPC. Disrupting this asymmetric centrosome segregation alters NPCs’ fate decisions and their maintenance in the Ventral Zone (VZ) of human cortical organoids (Royall *et al*., 2023). Many genes that regulate centrosome function and asymmetry have also been implicated in developmental disabilities, such as primary microcephaly (Nigg and Raff, 2009; Gilmore and Walsh, 2013; Conduit *et al*., 2015; Ramdas Nair *et al*., 2016; Link *et al*., 2019; Marthiens and Basto, 2020; Robinson *et al*., 2020; Jaiswal and Singh, 2021). However, there are fundamental gaps in our understanding of the underlying regulation of centrosome asymmetry, its function in ACD, and its impact on development.

*Drosophila* neural stem cells, called neuroblasts, provide an ideal model to study centrosome asymmetry. *Drosophila* neural stem cells divide asymmetrically, giving rise to a self-renewed neuroblast and a differentiating ganglion mother cell (GMC) (Gallaud *et al*., 2020). Neuroblasts invariably retain the daughter centriole-containing centrosome (hereafter daughter centrosome), whereas differentiating GMCs inherit the older mother centriole-containing centrosome (hereafter mother centrosome) (Rusan and Peifer, 2007; Homem and Knoblich, 2012; Januschke *et al*., 2013a; Lerit *et al*., 2013). These centrosomes further differ in MTOC activity, where the daughter centrosome maintains MTOC activity throughout interphase while the mother centrosome sheds PCM proteins in early interphase and remains inactive until centrosome maturation in prophase. Whereas microtubule-nucleating daughter centrosomes remain anchored to the apical cell cortex, the inactive mother centrosome drifts in the cytoplasm until prophase before being pulled to the basal cell cortex (Rebollo *et al*., 2007; Rusan and Peifer, 2007; Singh *et al*., 2014).

This difference in MTOC activity is determined by the centrosome’s molecular identity. In interphase neuroblasts, the apical centrosome retains the daughter-centriole protein Centrobin (Cnb) and the mitotic kinase Polo (Plk1). Polo and PCM proteins are shed from the basal centrosome via the actions of Pericentrin-like protein (Plp), Polo kinase 4 (Plk4) and Bld10 (Januschke *et al*., 2013a; Singh *et al*., 2014; Lerit *et al*., 2015; Ramdas Nair *et al*., 2016; Gambarotto *et al*., 2019; Gallaud *et al*., 2020). Centrobin interacts with *γ*Tubulin and other PCM members such as Plp, Sas-4, Sas-6, and Centrosomin (Cnn), and is required for the maintenance of MTOC activity in interphase (Januschke *et al*., 2013a; Ramdas Nair *et al*., 2016). Centrobin’s localization is dependent on phosphorylation by Polo, and proper microtubule nucleation is required for active recruitment of Polo to the apical centrosome (Januschke *et al*., 2013a; Gallaud *et al*., 2020). Recently, we showed that neuroblast centrioles replicate in mitosis, isochronous with a dynamic relocalization of Centrobin from the original mother centriole to the nascent daughter centriole. This transfer is facilitated by Polo-mediated phosphorylation of Cnb, such that by the end of mitosis, only the daughter centriole contains Cnb. Upon centriole separation, the younger Cnb^+^ centriole retains microtubule nucleating activity, tethering it to the apical neuroblast cortex, whereas the older Cnb^-^ centriole sheds Polo and PCM, thereby downregulating MTOC activity (Singh *et al*., 2014; Gallaud *et al*., 2020). Thus, dynamic Cnb relocalization in mitosis is necessary to establish centrosome asymmetry, manifested in biased MTOC activity in the following interphase (Gallaud *et al*., 2020).

The molecular mechanisms of centrosome asymmetry in general and Cnb relocalization in particular are largely unknown. Since Polo regulates Cnb‘s dynamic relocalization in mitosis, we reasoned that yet-to-be identified phosphatases could also regulate centrosome asymmetry in fly neural stem cells. Here, we characterize Protein phosphatase 4 (Pp4), a conserved serine/threonine phosphatase belonging to the phospho-protein phosphatase superfamily as a new regulator of neuroblast centrosome asymmetry *in vivo*. Pp4 regulates centrosome maturation, and spindle orientation in the developing neocortex and in stem cell development (Helps *et al*., 1998; Kloeker and Wadzinski, 1999; Sumiyoshi *et al*., 2002; Martin-Granados *et al*., 2008; Toyo-oka *et al*., 2008; Sousa-Nunes *et al*., 2009; Lyu *et al*., 2013; Voss *et al*., 2013; Xie *et al*., 2013; Karman *et al*., 2020a; Park and Lee, 2020). Pp4 localizes to centrosomes in both human neuronal progenitor cells and in developing fly embryos, where it is required for microtubule nucleation, growth, and stabilization (Helps *et al*., 1998). Pp4 forms a complex with two regulatory subunits Pp4R2 and Pp4R3 (Falafel, Flfl), the latter of which has been shown to be a key mediator of cell fate determinants in neuroblast divisions (Sousa-Nunes *et al*., 2009; Connell *et al*., 2021). We show that the removal of Pp4 results in a loss of MTOC activity in interphase neuroblasts without compromising centrosome maturation in mitosis. *Pp4* mutants also show altered Centrobin and Polo localization in interphase and mitosis. We propose that Pp4 plays dual roles in the regulation of centrosome asymmetry in neural stem cells: in interphase, Pp4 dephosphorylates *γ*Tubulin to promote microtubule nucleation, and in mitosis, Pp4 dephosphorylates Centrobin to facilitate its transfer from mother to daughter centriole.

## RESULTS

### Pp4 is required for centrosome MTOC asymmetry in neural stem cells

We identified Pp4-19C (Pp4, hereafter. PPP4C in humans) as a possible regulator of centrosome asymmetry in a phosphatase RNAi screen. To characterize Pp4’s role in centrosome asymmetry, we generated a CRISPR-Cas9 deletion allele (see methods), that removed *Pp4*’s entire coding sequence (*Pp4^Δ^* hereafter) and performed live cell imaging experiments in intact larval brains (see methods for details). To that end, we crossed the MTOC marker mCherry::Jupiter (Cabernard and Doe, 2009a) (Jupiter encodes a microtubule (MT)-binding protein (Karpova *et al*., 2006)) and the centriolar marker Asl::GFP (Blachon *et al*., 2008a) to *Pp4^Δ^* flies. As reported previously (Rebollo *et al*., 2007; Lerit *et al*., 2013; Singh *et al*., 2014; Ramdas Nair *et al*., 2016; Gallaud *et al*., 2020), the apical centrosome maintained robust MTOC activity in wild-type interphase neuroblasts, whereas the basal centriole contained little to no mCherry::Jupiter signal (Figure 1A; timepoints -00:32 and -00:29; orange and magenta dashed boxes, respectively; Movie 1). Upon entering prophase, the basal centrosome began to mature, forming an active MTOC (Figure 1A, timepoint -00:14; magenta dashed box). From prometaphase through metaphase, apical and basal centrosomes contained comparable levels of MTOC activity (timepoints -00:04 and 00:00, figure 1A, C, D). In contrast to wild type, *Pp4^Δ^* mutant neuroblasts failed to maintain an active apical MTOC in interphase (Figure 1B, timepoints -1:03 and - 0:36; Movie 02). However, both centrosomes matured normally and displayed robust mCherry::Jupiter signal from prometaphase onward (Figure 1B-D, timepoints -00:09 and -00:06).

**FIGURE 1:**
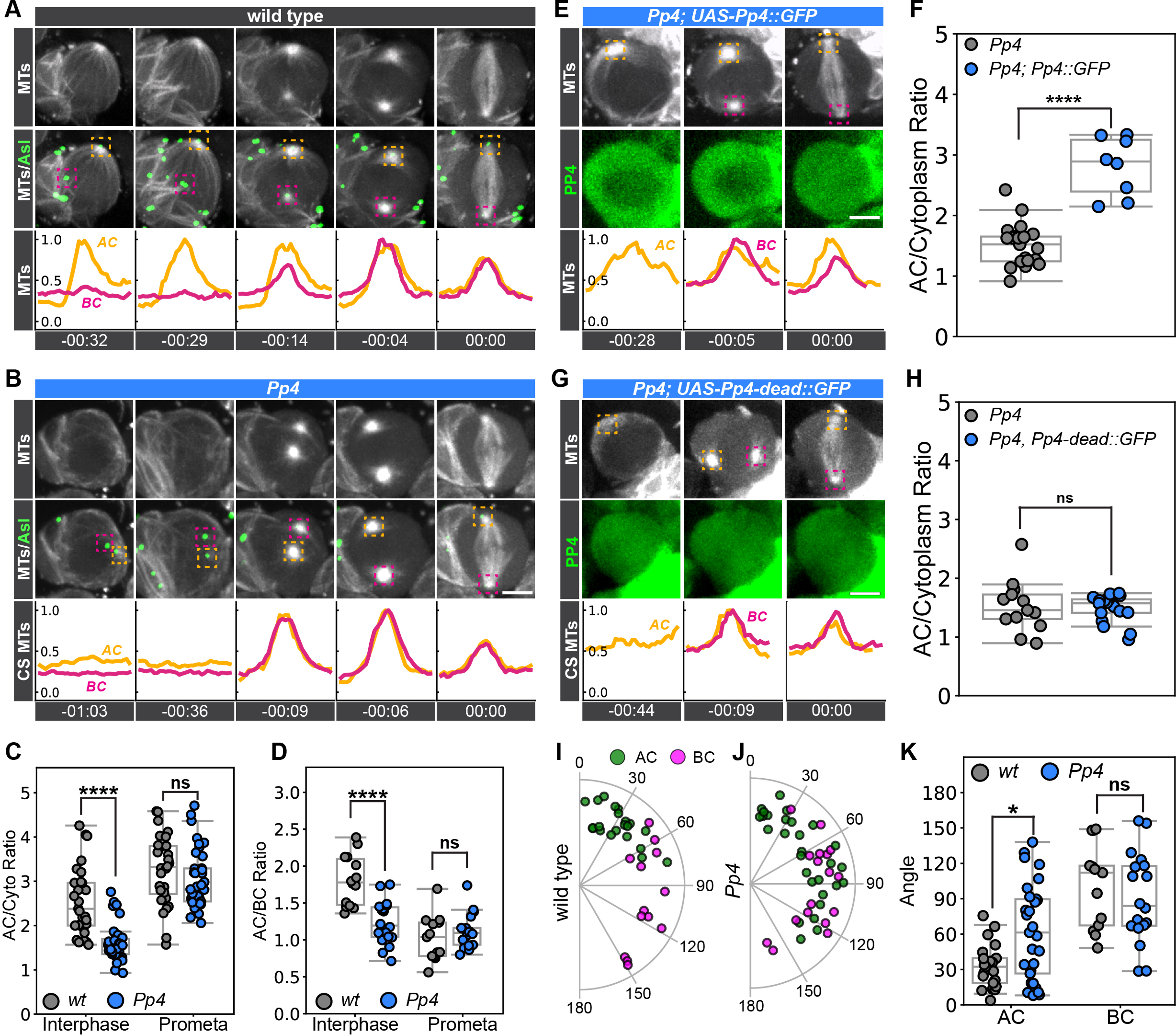
Pp4 is required for centrosome asymmetry in fly neural stem cells. Representative image sequences of a **(A)** wild type and **(B)** *Pp4^Δ^* mutant neuroblast, expressing *worGal4, UAS-mCherry::Jupiter* (grey, top row) and *Asterless::GFP* (green, merge on second row). Orange and magenta dashed boxes denote the apical (AC) and basal (BC) centrosome, respectively. Line scans of microtubule intensity at the apical (orange) and basal (magenta) centrosomes are shown below. **(C)** Normalized AC/Cytoplasm ratio of microtubule intensity in wild type (gray) and *Pp4^Δ^* mutant (blue) neuroblasts for interphase and prometaphase. **(D)** Normalized AC/BC ratio of microtubule intensity in wild type (gray) and *Pp4^Δ^* mutant (blue) neuroblasts for interphase and prometaphase. **(E)** Representative image sequence showing a *Pp4^Δ^* mutant neuroblast expressing *Worgal4, UAS-mCherry::Jupiter* (grey, top row) and *UAS-Pp4::GFP* (green, second row). **(F)** Normalized AC/Cytoplasm ratio of microtubule intensity in interphase *Pp4^Δ^* mutant neuroblasts (gray) and *Pp4^Δ^* mutant neuroblasts, expressing *UAS-Pp4::GFP* (blue) under control of *worGal4*. **(G)** Representative image sequence showing a *Pp4^Δ^* mutant neuroblast expressing *Worgal4, UAS-mCherry::Jupiter* (grey, top row) and *UAS-Pp4^[D85N, H115N]^::GFP* (green, second row). **(H)** Normalized AC/Cytoplasm ratio of microtubule intensity in interphase *Pp4^Δ^* (gray) and *Pp4^Δ^* mutant neuroblasts, expressing *UAS-Pp4^[D85N, H115N]^::GFP* (blue). Radial plots of apical (AC; green dots) and basal (BC; magenta dots) centrosome displacement between interphase and metaphase for **(I)** wild type and **(J)** *Pp4^Δ^* mutant neuroblasts. **(K)** Apical and basal centrosome displacement angles (between interphase and metaphase) are plotted for wild type (grey) and *Pp4^Δ^* mutant (blue) neuroblasts. Asl; Asterless. MTs; microtubules. Scale bar denotes 5 µm. Statistical significance between normal distributions of data was determined via Student’s unpaired t-test to compare the differences in the distribution of collected data. Significance between non-parametric distributions of data was determined with a two-sided Mann-Whitney U statistical test. *p <* 0.05 were considered significant; * *p <* 0.05, ** *p <* 0.01, *** *p <* 0.001, **** *p <* 0.0001. Time in: h:mins.

*Pp4^Δ^*’s loss of MTOC activity phenotype could be rescued by expressing full length wild type *Pp4* (5X-UAS-Pp4::GFP) in *Pp4^Δ^* mutants, using the neuroblast specific worniuGal4 (worGal4 (Albertson *et al*., 2004)) driver (Figure 1E, F). To test whether interphase MTOC activity depends on Pp4’s catalytic activity, we expressed a phosphatase-dead (10X-UAS-Pp4[D85N, H115N]) construct in *Pp4^Δ^* mutants. Pp4[D85N, H115N] carries mutations to the predicted active (D85) and binding (H115) sites of Pp4 (Umezawa *et al*., 2010; Park and Lee, 2020; Sandal *et al*., 2021). *Pp4^Δ^* mutant neuroblasts expressing 10X-UAS-Pp4[D85N, H115N] still lacked MTOC asymmetry, matching the phenotype seen in *Pp4* mutants (Figure 1G, H).

Sustaining an active MTOC in interphase allows the apical centrosome to maintain its position in the apical region, presumably via interactions with the cell cortex (Lerit *et al*., 2013; Singh *et al*., 2014; Ramdas Nair *et al*., 2016; Varadarajan and Rusan, 2018; Hannaford and Rusan, 2024). Since in *Pp4^Δ^* mutant neuroblasts the apical centrosome loses MTOC activity, we hypothesized that the lack of a robust interphase MTOC could compromise the centrosome’s apical tethering. To compare the apical centrosome’s position in wild type with *Pp4* mutants, we recorded the position of the centrosome in interphase and metaphase and calculated the deviation angle between the two time points. Apical wild type centrosomes do not move much between interphase and mitosis, consistent with an observed average angle of displacement of 33.0° (± 18.8°) (Figure 1I, K). Basal wild type centrosomes display more movement between interphase and metaphase, with an observed average displacement angle of 101° (± 36.0; Figure 1I, K). In *Pp4^Δ^* mutants, the average displacement angle of the apical centrosome – defined as the centrosome that segregates into the renewed neuroblast - increased to 55.7° (± 39.5), and the average displacement angle of the basal centrosome was 90.6° (± 38.2; Figure 1J, K). Comparing wild type with *Pp4^Δ^* mutant neuroblasts show that apical, but not basal centrosome positioning is more varied in *Pp4^Δ^* mutants (Figure 1K). Together, these data suggest that Pp4 maintains interphase MTOC activity in fly neural stem cells via its catalytic activity, which is necessary for tethering the active centrosome to the apical neuroblast cortex.

### Pp4’s regulatory subunits Pp4r2 and Pp4r3 are required for interphase MTOC activity

Pp4 is part of a heterotrimeric complex, consisting of the catalytic subunit Pp4c and two regulatory subunits, Pp4r2 and Pp4r3 (Pp4r3 is called Falafel (Flfl) in *Drosophila*) (Lipinszki *et al*., 2015a; Karman *et al*., 2020a; Park and Lee, 2020). Flfl is responsible for directing Pp4 to the centromere to regulate kinetochore integrity during mitosis and facilitates the basal localization of the basal cell fate determinant Miranda (Mira) (Sousa-Nunes *et al*., 2009; Lipinszki *et al*., 2015a). Additionally, Flfl physically interacts with the daughter-centriole protein Centrobin (Karman *et al*., 2020b). To test if Falafel is also required for interphase centrosome MTOC asymmetry, we imaged *flfl* mutant neuroblasts (*flfl[795]* (Sousa-Nunes *et al*., 2009) crossed to a deficiency, removing the entire *flfl* locus), expressing mCherry::Jupiter and Asl::GFP. Similar to *Pp4^Δ^* mutants, we observed that *flfl[795]/def* mutant neuroblasts lacked interphase MTOC activity (Supplemental Figure 1A-D).

Previous work identified Pp4r2 as a regulatory subunit responsible for binding to Pp4 substrates (Park and Lee, 2020). We sought to determine whether Pp4r2 was also required for regulating neuroblast centrosome asymmetry. We knocked-down Pp4r2 with inducible RNAi in fly neuroblasts, co-expressing the MTOC marker mCherry::Jupiter and the centriolar marker Asl::GFP. We observed that 25% of cells showed a phenotype identical to that seen in *Pp4^Δ^* mutants (timepoints 0:10 and 0:16 in Supplemental Figure 1E). We also observed a subset of cells that showed two active MTOCs in interphase (Supplemental Figure 1F-H). From these data, we conclude that both Pp4r2 and Pp4r3 are required for interphase MTOC asymmetry.

### Pp4 is required for interphase *γ*Tubulin localization at the apical centrosome

Pp4 interacts with *γ*Tubulin *in vitro* (Helps *et al*., 1998; Sumiyoshi *et al*., 2002; Martin-Granados *et al*., 2008; Alvarado-Kristensson *et al*., 2009; Voss *et al*., 2013; Karman *et al*., 2020a) and is required for centrosome maturation and microtubule regulation (Helps *et al*., 1998; Sumiyoshi *et al*., 2002; Martin-Granados *et al*., 2008). Further, phosphorylation of *γ*Tubulin has been shown to impact microtubule nucleation (Alvarado-Kristensson *et al*., 2009; Voss *et al*., 2013). Based on Pp4’s interactions with *γ*Tubulin, we hypothesized that Pp4 is required for proper *γ*Tubulin localization at centrosomes. There are two *γ*Tubulin isoforms in *Drosophila*: *γ*Tubulin23C, which has been shown to regulate centrosome activity in neuroblasts (Sunkel *et al*., 1995), and *γ*Tubulin37C, which controls meiosis during gametogenesis and early development (Tavosanis *et al*., 1997). We first sought to determine whether *γ*Tubulin37C and *γ*Tubulin23C are localized to centrosomes in both wild-type and *Pp4* mutant neuroblasts. In wild-type neuroblasts, expressing endogenously-tagged *γ*Tubulin23C::GFP (Mukherjee *et al*., 2020), the apical centrosome contains robust levels of *γ*Tubulin23C throughout interphase, while the basal centrosome only recruits *γ*Tubulin23C upon entry into mitosis. During mitosis, *γ*Tubulin23C localizes on both centrosomes (Figure 2A, C; Movie 3). In *Pp4^Δ^* mutants, we observed that *γ*Tubulin23C levels are reduced by ∼ 50% on the apical interphase centrosome compared to wild type but showed similar levels at prometaphase (Figure 2B, C; Movie 4).

**FIGURE 2:**
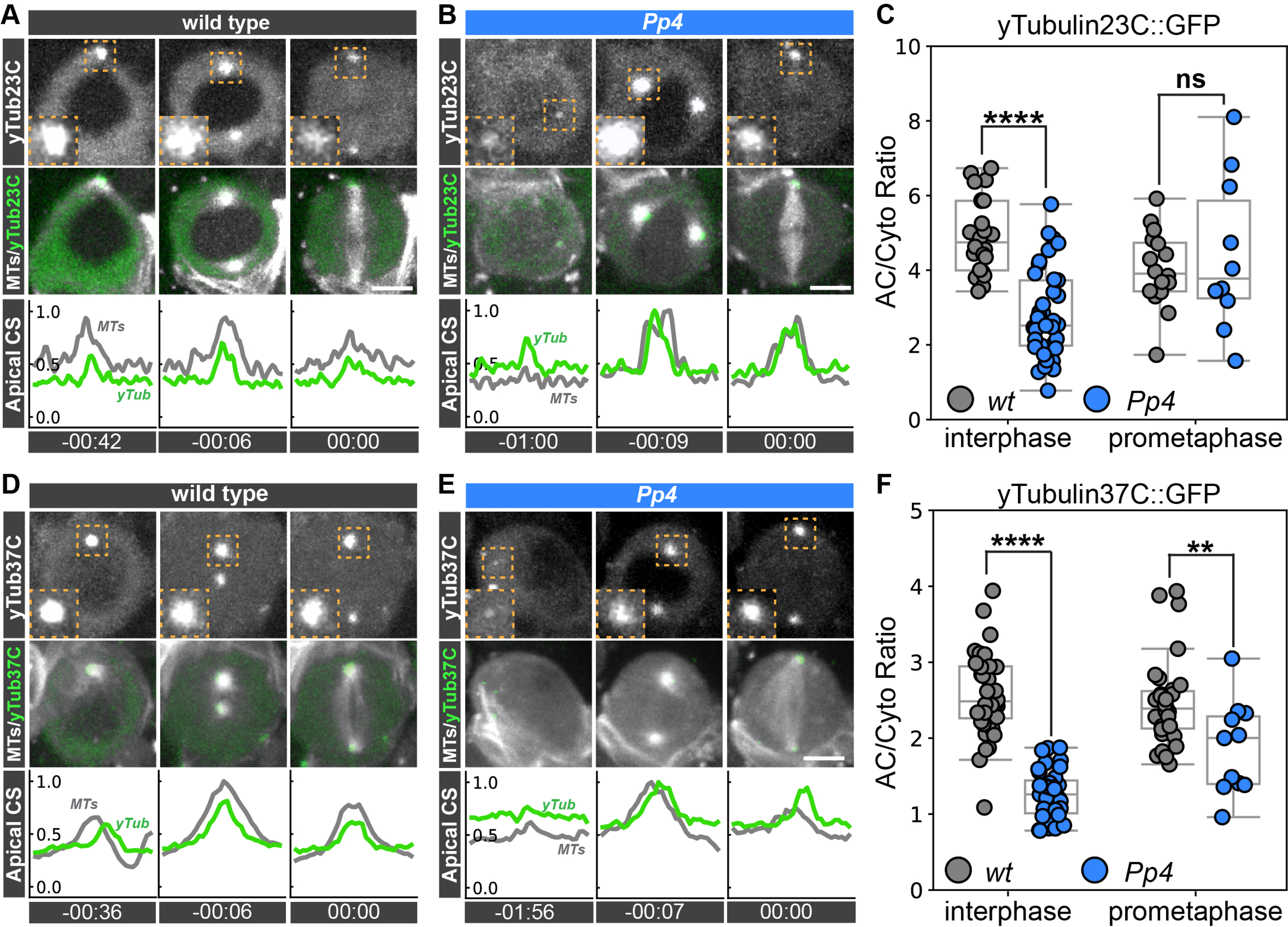
Pp4 is required for *γ*Tubulin localization at the apical centrosome in interphase. Representative image sequences of a **(A)** wild type or **(B)** *Pp4^Δ^* mutant neuroblast expressing *worGal4, UAS-mCherry::Jupiter* and **γ*Tubulin23C::GFP*. *γ*Tubulin23C::GFP (top row: white; middle row; green) and mCherry::Jupiter (middle row: white). Inserts in the top row show high magnification images of the apical centrosome (dashed orange box). Bottom row shows normalized microtubule (MTs; grey line) and *γ*Tubulin23C (γTub; green line) intensity line scans for the apical centrosome. **(C)** Quantification of *γ*Tubulin23C intensity at apical centrosomes, normalized to cytoplasmic signal, in interphase and prometaphase for wild type (gray) and *Pp4^Δ^* mutant (blue) neuroblasts. Representative image sequence of a **(D)** wild type or **(E)** *Pp4^Δ^* mutant neuroblast expressing *worGal4, UAS-mCherry::Jupiter* and *ncd-*γ*Tubulin37C::GFP*. ncd-*γ*Tubulin37C::GFP (top row: white; middle row; green) and mCherry::Jupiter (middle row: white). Inserts in the top row show high magnification images of the apical centrosome (dashed orange box). Bottom row shows normalized microtubule (MTs; grey line) and *γ*Tubulin37C (*γ*Tub; green line) intensity line scans for the apical centrosome. **(F)** Quantification of *γ*Tubulin23C intensity at apical centrosomes, normalized to cytoplasmic signal, in interphase and prometaphase for wild type (gray) and *Pp4* mutant (blue) neuroblasts. MTs; microtubules. CS; centrosome. Scale bar denotes 5 µm. Statistical significance between normal distributions of data was determined via Student’s unpaired t-test to compare the differences in the distribution of collected data. Significance between non-parametric distributions of data was determined with a two-sided Mann-Whitney U statistical test. *p <* 0.05 were considered significant; * *p <* 0.05, ** *p <* 0.01, *** *p <* 0.001, **** *p <* 0.0001. Time in: h:mins.

In wild-type cells, *γ*Tubulin37C::GFP expressed under the ncd promoter (Hallen *et al*., 2008b, 2008a), displayed robust localization at the apical centrosome in interphase. The basal centrosome lacked *γ*Tubulin37C in interphase but later acquired *γ*Tubulin37C upon entry into mitosis (Figure 2D, F). In contrast to this, *Pp4* mutants displayed low levels of *γ*Tubulin37C in interphase. Upon entry into mitosis, *γ*Tubulin37C localized to both centrosomes in *Pp4* mutants, albeit at significantly lower levels compared to wild type (Figure 2E, F). These data suggest that Pp4 is required for the proper localization of *γ*Tubulin23C and *γ*Tubulin37C at centrosomes in interphase neuroblasts.

### *γ*Tubulin37C and *γ*Tubulin23C mutants disrupt centrosome asymmetry

*γ*Tubulin contributes to the formation of *γ*Tubulin Ring Complexes (*γ*TuRCs), which promotes microtubule nucleation. These *γ*TuRCs are rapidly incorporated into the PCM of mature centrosomes, where they contribute to the formation of active MTOCs (Nigg and Stearns, 2011; Böhler *et al*., 2021). Both *γ*Tubulin23C and *γ*Tubulin37C are required for *γ*TuRC integrity in flies (Böhler *et al*., 2021). To clarify the role of both *γ*Tubulin isoforms in interphase MTOC formation in fly neural stem cells, we live-imaged mutant alleles for *γ*Tubulin23C (*γ*Tubulin23C^[A14-9]^, *γ*Tubulin23C^[A9-2]^, and *γ*Tubulin23C^[A15-2]^) and *γ*Tubulin37C (*γ*Tubulin37C^[3]^ and *γ*Tubulin37C^[e00793]^) with mCherry::Jupiter and Sas4::GFP and assessed MTOC activity in interphase (Wilson and Borisy, 1998a; Vázquez *et al*., 2008; Hughes *et al*., 2011). Several *γ*Tubulin alleles showed altered MTOC asymmetry: 7% of *γ*Tubulin23C^[A6-2]^, 74% of *γ*Tubulin23C^[A14-9]^ and 54% of *γ*Tubulin23C^[A15-2]^ mutant neuroblasts displayed two interphase MTOCs. Similarly, 70% of *γ*Tubulin37C^[e00793]^ and 42% of *γ*Tubulin37C^[3]^ mutants contained two interphase MTOCs (Supplemental Figure 2A-E). However, 20% of *γ*Tubulin37C^[3]^ mutants lacked discernable MTOC activity in interphase, the same phenotype as *Pp4^Δ^* mutants, albeit less penetrant (Figure 3A – F, Supplemental Figure 2E). To determine remaining protein levels in *γ*Tubulin23C and *γ*Tubulin37C mutants, we stained *γ*Tubulin23C^[A14-9]^ and *γ*Tubulin37C^[3]^ mutant larval brains with an anti-*γ*Tubulin antibody that recognizes both isoforms of fly *γ*Tubulin (see methods for details). Neither *γ*Tubulin37C^[3]^ nor *γ*Tubulin23C^[A14-9]^ mutants showed a loss of *γ*Tubulin at centrosomes in interphase or metaphase, respectively (Supplemental Figure 3A-D). Additionally, both *γ*Tubulin37C^[3]^ and *γ*Tubulin23C^[A14-9]^ showed increased presence of *γ*Tubulin on the basal centrosome, which is consistent with the observation of two active MTOCs in our live-cell data (Supplemental Figure 3A-D). For these experiments, *γ*Tubulin23C^[A14-9]^ and *γ*Tubulin37C^[3]^ were crossed over deficiency chromosomes, removing the entire *γ*Tubulin23C or *γ*Tubulin37C coding region, respectively. Since these crosses create strong loss-of-function conditions, it is unlikely that the presence of *γ*Tub is a consequence of low allele penetrance. Considering that *γ*Tubulin37C and *γ*Tubulin23C both contribute to the formation of a *γ*TuRC (Tariq *et al*., 2020), it is more likely that the two isoforms are redundant with each other. To test for genetic redundancy, we created a recombinant *γ*Tubulin37C^[3]^, *γ*Tubulin23C^[A15-2]^ chromosome. *γ*Tubulin37C^[3]^, *γ*Tubulin23C^[A15-2]^ double-mutant interphase and mitotic neuroblasts centrosomes were devoid of *γ*Tubulin in second instar larval brains (they did not mature into third instar larvae; Supplemental Figure 3E, F). These data suggest that both *γ*Tubulin37C and *γ*Tubulin23C are required for MTOC asymmetry in interphase neural stem cells.

**FIGURE 3:**
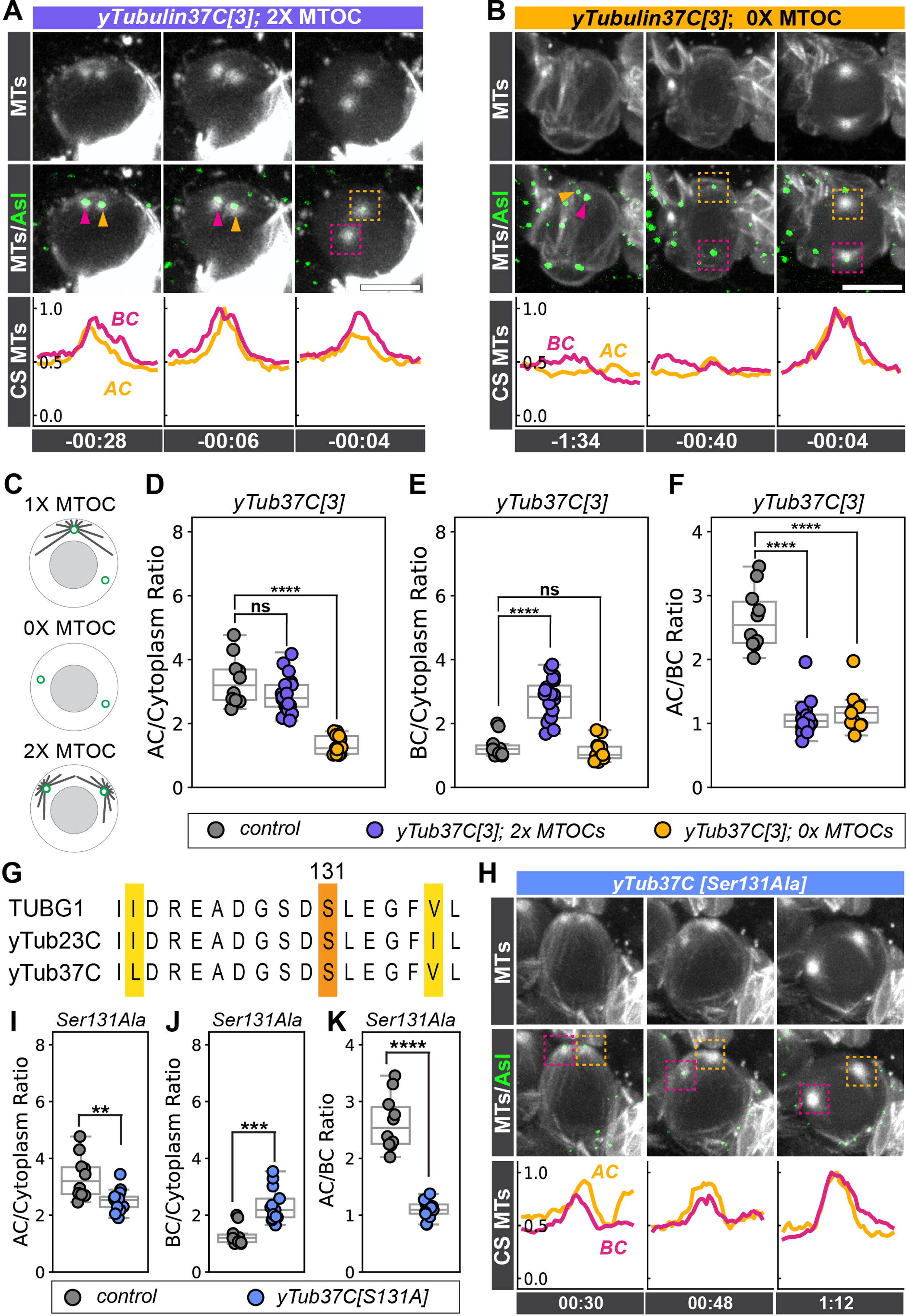
*γ*Tubulin mutants exhibit impaired interphase centrosome asymmetry. Representative time series of a **(A)** **γ*Tubulin37C^[3]^* mutant neuroblast expressing *worGal4, UAS-mCherry::Jupiter* (white, top row) and Sas4::GFP (green, merge in second row). Apical (AC; magenta) and basal centrosomes (BC; orange) are highlighted with magenta and orange arrowheads and dashed boxes, respectively. Microtubule (MT) intensity line scans of the apical (orange) and basal (magenta) centrosome (CS) is shown below. Note that the shown *γ*Tubulin37C^[3]^ mutant neuroblast has two active MTOCs (2xMTOC). **(B)** Representative *γ*Tubulin37C^[3]^ mutant neuroblast with no active MTOC (0x MTOC). **(C)** Schematic illustration of the different MTOC phenotypes. **(D)** Normalized AC/Cytoplasm, **(E)** BC/Cytoplasm, or **(F)** AC/BC ratios for wild type (grey circles) and *γ*Tubulin37C^[3]^ mutants (purple circles: 2xMTOC phenotype; yellow circles: 0xMTOC phenotype). **(G)** Protein sequence alignment of human TUBG1, *γ*Tubulin23C, and *γ*Tubulin37C. Variations in sequence are highlighted in yellow. The conserved Serine at position 131 is highlighted in orange. **(H)** Representative image sequence of a *γ*Tubulin37C^[S131A]^ mutant neuroblast, expressing worGal4, UAS-mCherry::Jupiter (white, top row) and Sas4::GFP (green, merge in second row). Normalized **(I)** AC/Cytoplasm, **(J)** BC/Cytoplasm, or **(K)** AC/BC ratios in wild type controls (gray) and *γ*Tubulin37C^[S131A]^ mutants (blue). MTs; microtubules. CS; centrosome. Scale bar denotes 5 µm. Statistical significance between normal distributions of data was determined via Student’s unpaired t-test to compare the differences in the distribution of collected data. Significance between non-parametric distributions of data was determined with a two-sided Mann-Whitney U statistical test. *p <* 0.05 were considered significant; * *p <* 0.05, ** *p <* 0.01, *** *p <* 0.001, **** *p <* 0.0001. Time in: h:mins.

### *γ*Tubulin phospho-mutants exhibit impaired centrosome asymmetry

In U2OS human cell culture, phosphorylation of *γ*Tubulin at Serine 131 prevents microtubule nucleation, implicating phosphorylation of Serine 131 as a regulator of MTOC formation (Alvarado-Kristensson *et al*., 2009). Both *Drosophila γ*Tubulin isoforms are highly conserved with Human TUBG1 at and around Serine 131 (Figure 3G). We therefore hypothesized that Pp4-mediated dephosphorylation of *γ*Tubulin at Serine 131 could promote microtubule nucleation. Because *γ*Tubulin37C^[3]^ mutants exhibited a loss of function phenotype similar to the *Pp4^Δ^* phenotype, we chose to examine the requirement of Serine 131 phosphorylation of *γ*Tubulin37C. To test this hypothesis, we mutated endogenous Serine 131 in *γ*Tubulin37C to Alanine (*γ*Tubulin^[S131A]^) using CRISPR/Cas9 (see methods) and crossed this allele to a deficiency stock that lacks the entire *γ*Tubulin37C coding region. We reasoned that the S131A mutant would mimic the unphosphorylated state of *γ*Tubulin37C and display an increase in MTOC activity in interphase neuroblasts. Indeed, 60% of *γ*Tubulin^[S131A]^ neuroblasts displayed two active interphase MTOCs, albeit with lower mCherry::Jupiter levels compared to wild type (Figure 3H-K; Supplemental Figure 2E). *γ*Tubulin^[S131A]^ mutants also showed elevated levels of *γ*Tubulin at the basal centrosome in interphase, consistent with the observation of two interphase MTOCs in our *in vivo* movies (Supplemental Figure 3A-D). These data suggests that the phosphorylation state of *γ*Tubulin at Serine 131 is involved in the maintenance of interphase MTOCs in *Drosophila* neuroblasts.

### Cdk1 contributes to interphase centrosome asymmetry

*In vitro,* Cyclin-dependent kinase 1 (Cdk1) phosphorylates the *γ*Tubulin residues Serine 80 and Threonine 196. Cdk1 also directly phosphorylates PP4r2 and PP4r3, which is predicted to decrease the catalytic activity of the Pp4 complex (Toyo-oka *et al*., 2008; Voss *et al*., 2013). Therefore, Cdk1 could regulate *γ*Tubulin via direct phosphorylation, or indirectly by inactivating Pp4 (Figure 4A). Phosphorylated *γ*Tubulin is predicted to result in reduced microtubule nucleation, as suggested previously (Alvarado-Kristensson *et al*., 2009; Voss *et al*., 2013). To further explore the role of *γ*Tubulin regulation via phosphorylation, we first knocked down Cdk1 in larval neuroblasts using the neuroblast-specific driver worGal4 and imaged MTOC activity with mCherry::Jupiter. In either model, the removal of Cdk1 would decrease *γ*Tubulin phosphorylation, potentially increasing microtubule nucleation in interphase. Indeed, in contrast to wild-type neuroblasts, we observed that 38% of Cdk1 RNAi-expressing neuroblasts display interphase MTOC activity on both centrosomes (Figure 4B, F-H; timepoints -1:18). However, in 24% of Cdk1 RNAi expressing neuroblasts, neither centrosome displayed interphase MTOC activity, similar to *Pp4^Δ^* mutants (Figure 4C, F-H; timepoints -1:24). The former finding suggests that Cdk1 has an inactivating role in microtubule nucleation, whereas the latter phenotype is more in line with a model where Cdk1 has a microtubule-nucleation-promoting function. If Cdk1 downregulates MTOC activity, an increase in Cdk1 levels on neuroblast centrosomes should cause a decrease in microtubule nucleation, manifested in a Pp4-like MTOC phenotype. To this end, we employed a previously used nanobody strategy, localizing the single-chain anti-GFP nanobody (vhhGFP4; (Saerens *et al*., 2005; Caussinus *et al*., 2012)) to centrosomes with the PACT domain (Gillingham and Munro, 2000; Gallaud *et al*., 2020). Crossing this PACT-vhhGFP4 construct to endogenously-tagged Cdk1::GFP should enrich Cdk1 at neuroblast centrosomes throughout the cell cycle. In wild type neuroblasts, Cdk1::GFP localizes to the apical centrosome in interphase and appears on both centrosomes in early mitosis (Figure 4D). However, crossing Cdk1::GFP with the PACT-vhhGFP4 construct enriched Cdk1 on centrosomes in interphase and mitosis. Under these conditions, 60% of interphase neuroblasts displayed no MTOC activity, similar to *Pp4^Δ^* mutants. However, mitotic spindle formation progressed normally (Figure 4E-H). Although the exact mechanisms are unknown, these data suggest that Cdk1 is required for regulating centrosome asymmetry in interphase neuroblasts.

**FIGURE 4:**
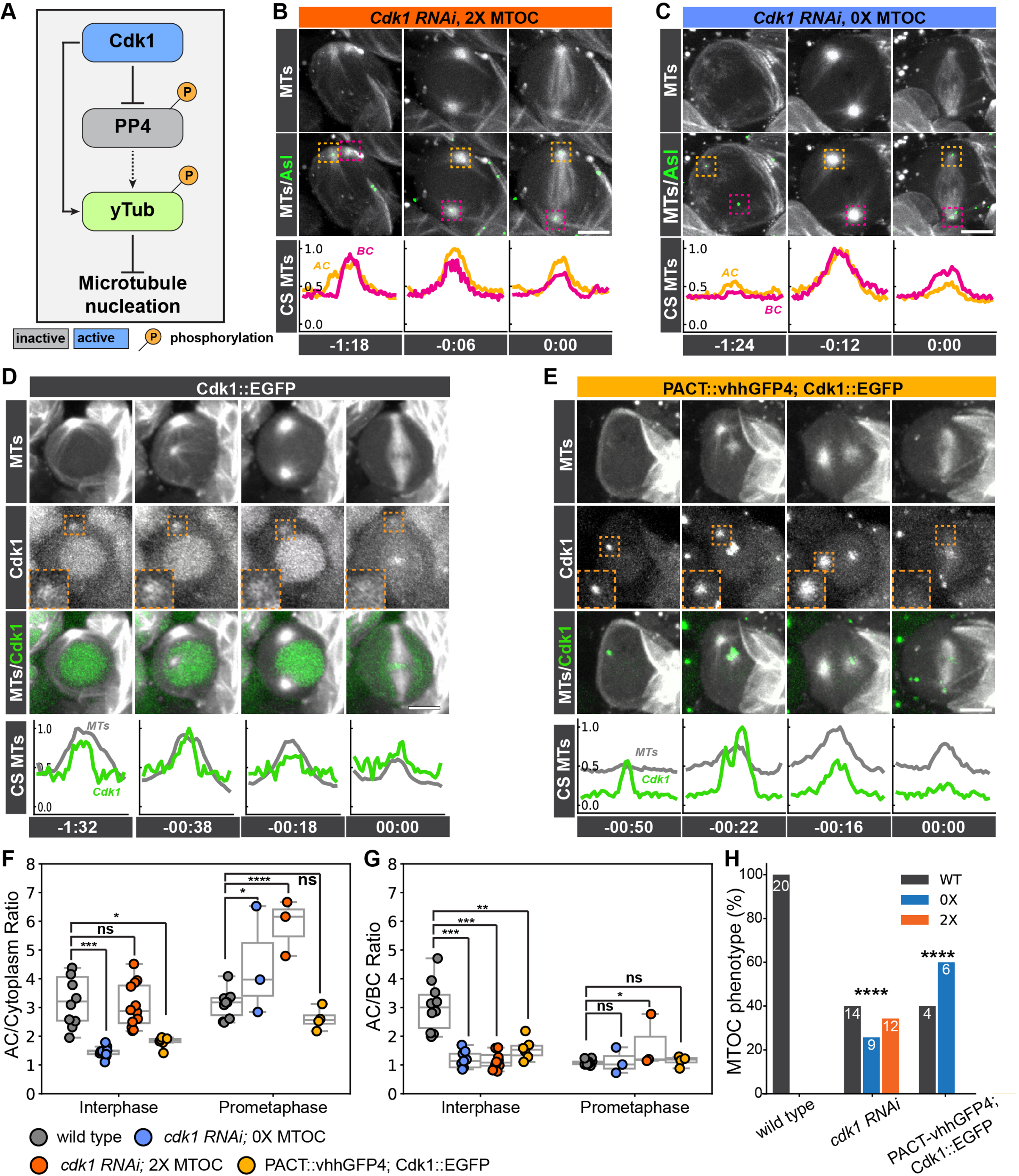
Cdk1 is required for centrosome asymmetry. **(A)** Working model from (Alvarado-Kristensson *et al*., 2009; Voss *et al*., 2013): active Cdk1 could inhibit Pp4 via phosphorylation or Cdk1 could directly phosphorylate γTub. Phosphorylated γTub is proposed to decrease microtubule nucleation activity. **(B, C)** Representative image sequences of *cdk1 RNAi*, worGal4, UAS-mCherry::Jupiter (top row: white), and Asl::EGFP (middle row: green) expressing neuroblasts showing **(B)** a 2xMTOC or **(C)** 0xMTOC phenotype. Orange and magenta dashed boxes highlight apical (AC) and basal (BC) centrosomes, respectively. Bottom row shows normalized line scans of microtubule intensity at the apical (yellow line) and basal (magenta line) centrosome. Representative time series of a **(D)** wild type neuroblast, expressing *Cdk1::EGFP* (middle row: white; green in merge) and *worGal4 UAS-mCherry::Jupiter* (top row and merge: white). **(E)** Representative neuroblast expressing, worGal4 UAS-mCherry::Jupiter (top row and merge: white), *Cdk1::EGFP* (middle row: white; green in merge) and the nanobody UAS-PACT::*vhhGFP4*. Insets show high magnification images of Cdk1::EGFP signal at the apical centrosome. Quantification of **(F)** AC/Cytoplasm or **(G)** AC/BC ratios in wild type (gray), *cdk1* RNAi (blue and orange circles; 0xMTOC and 2xMTOC, respectively) and nanobody expressing (*PACT::vhhGFP4; Cdk1::EGFP;* yellow circles) neuroblasts. **(H)** Frequency of 1X, 2X, or 0X MTOCs in wild type, *cdk1* RNAi, and Cdk1 trapping experiments. Numbers on bars indicate the number of measured neuroblasts. Asl; Asterless. MTs; microtubules. Scale bar denotes 5 µm. Statistical significance between normal distributions of data was determined via Student’s unpaired t-test to compare the differences in the distribution of collected data. Significance between non-parametric distributions of data was determined with a two-sided Mann-Whitney U statistical test. The chi-squared goodness-of-fit statistical test was done to test for differences between frequencies of observed phenotypes. *p <* 0.05 were considered significant; * *p <* 0.05, ** *p <* 0.01, *** *p <* 0.001, **** *p <* 0.0001. Time in: h:mins.

### Pp4 is required for apical centrosome localization of Polo but not Centrobin in interphase

We and others previously showed that centrosome asymmetry requires the mitotic kinase Polo (Plk1 in vertebrates) and the daughter-centriole protein Centrobin (Cnb; CNTROB in humans). Reducing Polo or loss of Cnb causes loss of apical MTOC activity in interphase neuroblasts, similar to the *Pp4* mutant phenotype described here (Januschke *et al*., 2013a; Singh *et al*., 2014; Ramdas Nair *et al*., 2016; Gallaud *et al*., 2020). We therefore sought to determine whether Pp4 contributes to the localization of either Polo or Cnb during interphase. We imaged *Pp4^Δ^* mutant larval neuroblasts, expressing the MTOC marker mCherry::Jupiter in conjunction with either Polo::EGFP or Cnb::EGFP (Gallaud *et al*., 2020). In wild-type neuroblasts, Polo is localized on the apical centrosome in interphase (Figure 5A, C; timepoints -01:18 and -00:24). Upon entry into mitosis, a subset of Polo is localized to kinetochores, while another fraction is retained at both the apical and basal centrosomes (Figure 5A, C; timepoints 00:00). In contrast, *Pp4* mutant neuroblasts showed significantly reduced Polo localization at the apical centrosome in interphase, and a significant increase in Polo localization during centrosome maturation (Figure 5B, C; timepoints -01:00 and 00:00). Additionally, Polo is retained on centrosomes in mitosis but was not detected on kinetochores in *Pp4* mutants (Figure 5B; timepoints 00:00). To better quantify the distribution of Polo in *Pp4* mutants, we imaged Polo::EGFP together with Asl::mCherry (Conduit *et al*., 2015) and measured Polo’s distribution with line scan measurements (Figure 5D, E; see methods for details). In contrast to wild-type centrosomes, which display a sharp peak centered on Asl, Polo is more diffusely localized in *Pp4^Δ^* mutant neuroblasts (Figure 5F, G).

**FIGURE 5:**
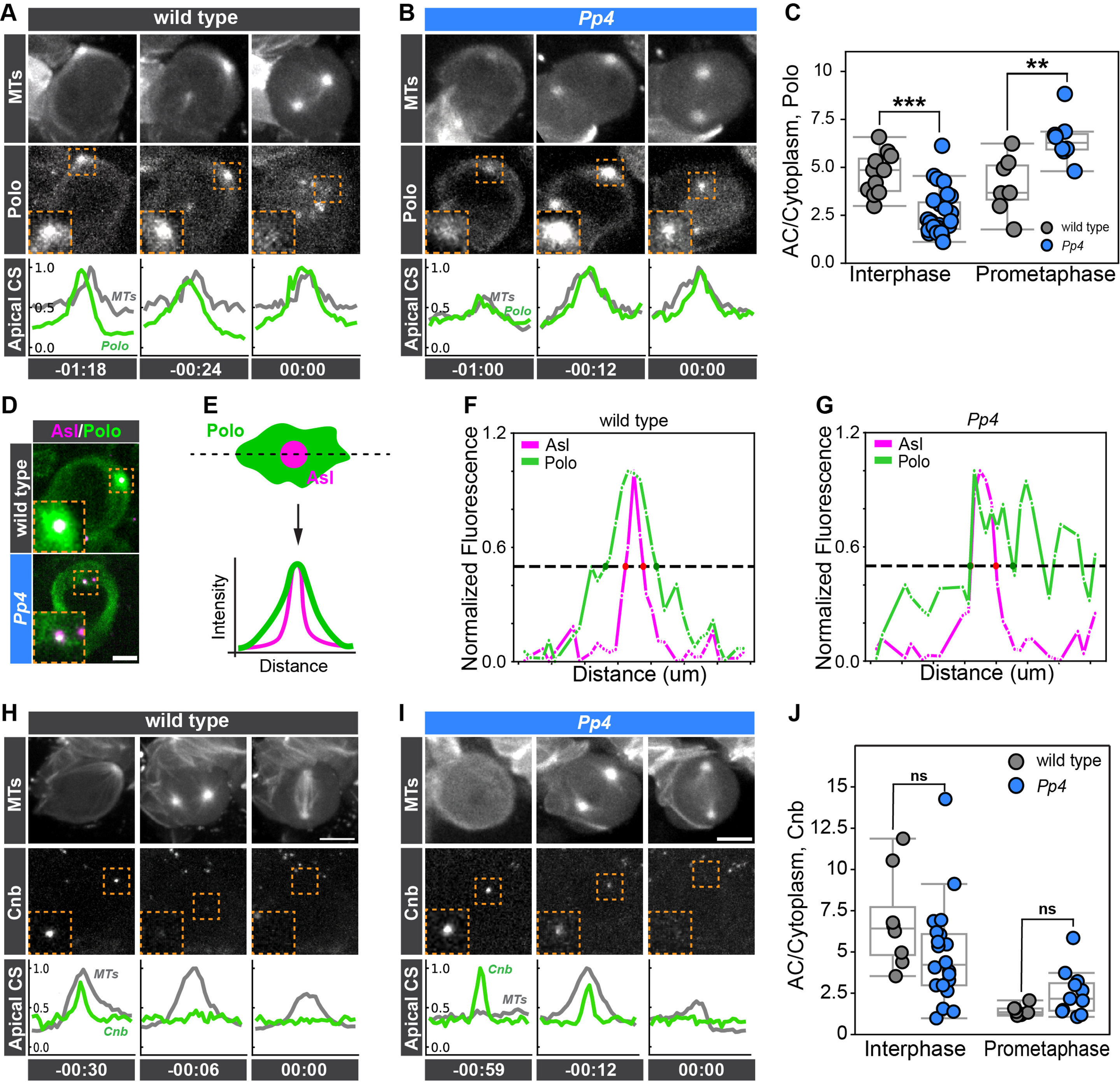
Pp4 is required for robust Polo but not Centrobin localization on the apical interphase centrosome. Representative image sequences of a **(A)** wild-type or **(B)** *Pp4^Δ^* mutant neuroblast expressing *worGal4, UAS-mCherry::Jupiter* (top row: white), *Polo::GFP* (middle row: white). Orange dashed boxes highlight apical centrosomes. High magnification images are shown in the bottom left corner. Normalized microtubule (gray line) and Polo::GFP (green line) intensity line scans for the apical centrosome are shown below. **(C)** Quantification of AC/Cytoplasm ratio for Polo::GFP intensity in wild type (gray), and *Pp4^Δ^* mutants (blue) at interphase and prometaphase. **(D)** Representative Images of a wild type (top row) or *Pp4^Δ^* mutant (bottom row) neuroblast, expressing endogenous Polo::EGFP (green) and Asterless::mCherry (magenta). **(E)** Schematic and representative intensity line scans to quantify Polo distribution at the centrosome are shown for **(F)** wild type and **(G)** *Pp4^Δ^* mutants. The line scans correspond to the cells shown in D. Green and magenta lines represent Polo and Asl intensity profiles. The black dashed line indicates the width at half height for each distribution. Representative image sequence of a **(H)** wild type or **(I)** *Pp4* mutant neuroblast expressing *worGal4, UAS-mCherry::Jupiter* (top row: white) and *Centrobin::GFP (*middle row: white). Corresponding intensity line scans are shown below. **(J)** Quantification of AC/Cytoplasm ratio of Centrobin::GFP fluorescence intensity in wild type (gray), and *Pp4^Δ^* mutants (blue) at interphase and prometaphase. Asl; Asterless. MTs; microtubules. Scale bar denotes 5 µm. Statistical significance between normal distributions of data was determined via Student’s unpaired t-test to compare the differences in the distribution of collected data. Significance between non-parametric distributions of data was determined with a two-sided Mann-Whitney U statistical test. *p <* 0.05 were considered significant; * *p <* 0.05, ** *p <* 0.01, *** *p <* 0.001, **** *p <* 0.0001. Time in: h:mins.

Centrobin is a substrate of Polo and needs to be phosphorylated by Polo for its centrosomal localization (Januschke *et al*., 2013a). In wild-type interphase neuroblasts, Centrobin is localized only to the apical daughter centrosome in interphase and weakly localized on both centrosomes in mitosis (Figure 5H and (Gallaud *et al*., 2020)). Surprisingly, in *Pp4^Δ^* mutants, Cnb is normally localized in interphase centrosomes but slightly elevated in prometaphase (Figure 5I, J). From these data we conclude that Pp4 is required for robust and confined Polo, but not Cnb localization at the apical interphase centrosome.

### Pp4 is required for Centrobin’s enrichment on the daughter centriole during mitosis

Asymmetric MTOC activity, manifested in one active and one inactive interphase MTOC, is regulated two-fold during the neuroblast cell cycle: (1) in mitosis, Centrobin relocalizes from the mother to the daughter centriole via downregulation of Centrobin on the mother centriole. This is followed by a transfer of the remaining Centrobin, originating from the mother centriole to the daughter centriole. Subsequently, Centrobin increases on the daughter centriole in late mitosis and early interphase. This dynamic Centrobin localization establishes two molecularly distinct centrosomes, which is necessary for interphase MTOC activity (Gallaud *et al*., 2020). (2) As neuroblasts enter interphase after cytokinesis, PCM components are shed from the Cnb^-^ mother centrosome that is destined to segregate into the differentiating GMC. PCM is retained on the Cnb^+^ daughter centrosome, which remains in the neuroblast (Rebollo *et al*., 2007; Lerit *et al*., 2013; Singh *et al*., 2014). We therefore hypothesized that in addition to Polo, Pp4 could be required for the localization of Centrobin onto the daughter centriole during mitosis. To test this, we performed three-dimensional structured illumination microscopy (3D-SIM) imaging experiments of control and *Pp4^Δ^* mutant neuroblasts expressing Cnb::EGFP. We used microtubules as a reference marker for cell cycle progression and Asterless to differentiate between the mother and daughter centrioles. As previously shown, Asterless intensity increases with centriolar age, providing a Centrobin-independent marker to distinguish between mother and daughter centrioles (Novak *et al*., 2014; Fu *et al*., 2016; Gallaud *et al*., 2020). We measured Centrobin levels on the mother and daughter centriole and plotted the corresponding daughter-to-mother intensity ratios to quantify the difference between control and *Pp4^Δ^* mutant neuroblasts (Figure 6D, E, Supplemental Figure 4A). As previously shown, in control neuroblasts, Centrobin is first localized on the mother centriole in prophase and prometaphase before enriching on the daughter centriole during metaphase, anaphase and telophase (Figure 6A, D, E; Supplemental Figure 4B-E). In all control neuroblasts, only the daughter centriole contained Cnb in telophase (Figure 6A, D-H). In *Pp4^Δ^* mutants, we observed mother centrioles with Centrobin in prophase and prometaphase, comparable to wild type neuroblasts (Figure 6B, D, E). However, from metaphase through telophase, we observed three distinct phenotypes, differing from wild type neuroblasts: (1) Cnb was either completely absent, (2) incorrectly enriched on the mother centriole, or (3) present on both the mother and daughter centriole (Figure 6B, E-H).

**FIGURE 6:**
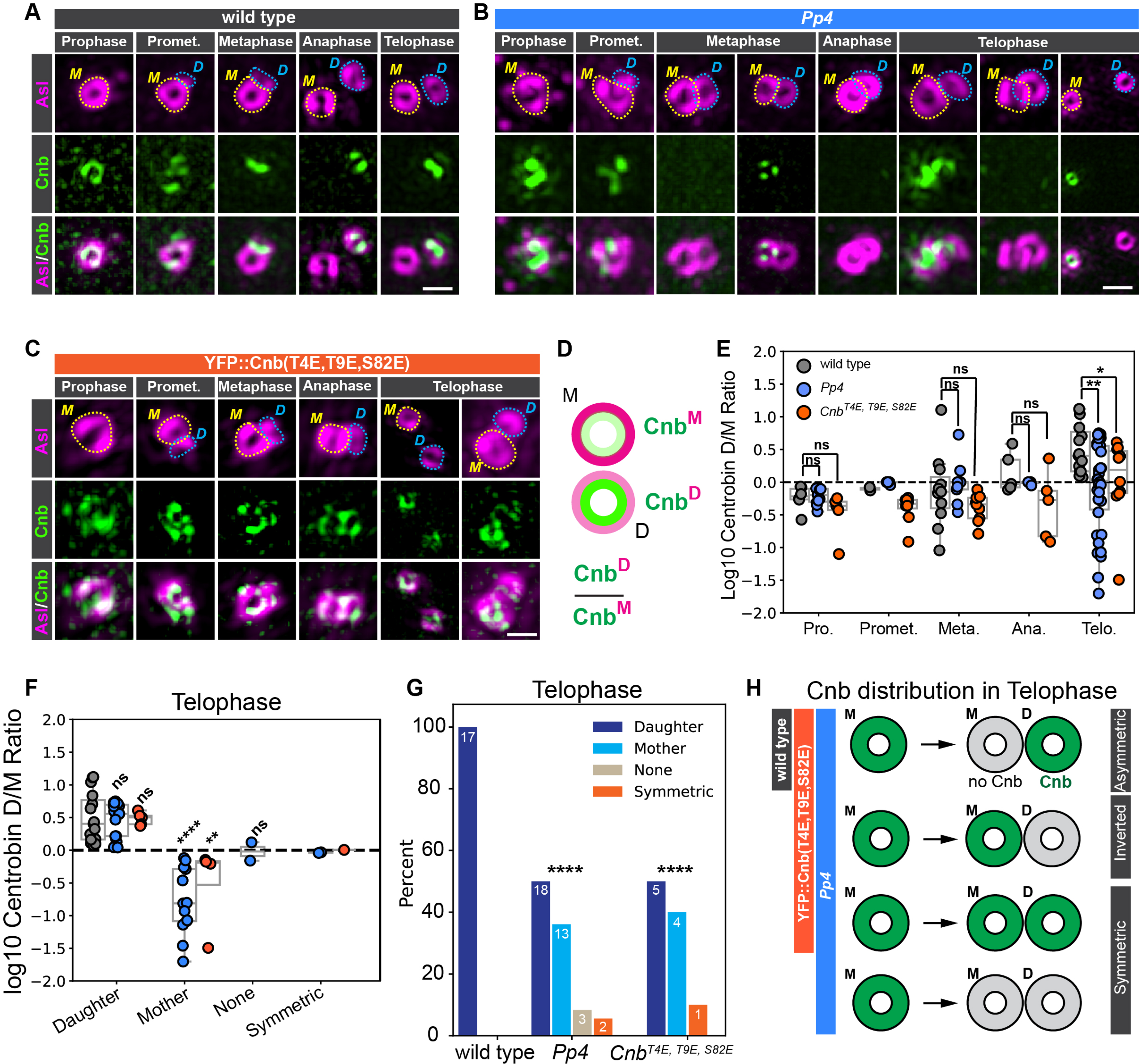
Pp4 is required for Centrobin localization in mitosis. Representative 3D-SIM images for **(A)** Cnb::EGFP expressing wild type, **(B)** *Pp4^Δ^* or **(C)** YFP::Cnb^T4E,T9E,S82E^ expressing neuroblasts. All samples were co-stained with anti-Asterless (Asl; top row; magenta). Middle row shows Cnb::EGFP (A, B) or YFP::Cnb^T4E,T9E,S82E^ (C). Merged images are shown in the bottom row. Mother and daughter centrioles are outlined with yellow and blue dashed circles, respectively. **(D)** Mother centrioles have higher Asl intensity. Centrobin localization at distinct mitotic stages was determined by calculating the ratio of daughter-to-mother centriole Cnb. **(E)** Centrobin D/M Log_10_-normalized ratios for prometaphase, metaphase, anaphase, and telophase for wild type (gray circles), *Pp4^Δ^* (blue circles) and YFP::Cnb^T4E,T9E,S82E^ expressing neuroblasts (orange circles). **(F)** Centrobin D/M Log_10_-normalized telophase enrichment ratios and **(G)** corresponding bar graphs for wild type (gray circles), *Pp4^Δ^* (blue circles) and YFP::Cnb^T4E,T9E,S82E^ (orange circles), separated by enrichment. Numbers on bars denote number of measured neuroblasts. **(H)** Schematic summary of telophase Cnb localization in the wild type, *Pp4^Δ^* and YFP::Cnb^T4E,T9E,S82E^ expressing neuroblasts. Asl; Asterless. Cnb; Centrobin. Scale bar denotes 0.5 µm. Statistical significance between normal distributions of data was determined via Student’s unpaired t-test to compare the differences in the distribution of collected data. Significance between non-parametric distributions of data was determined with a two-sided Mann-Whitney U statistical test. The chi-squared goodness-of-fit statistical test was done to test for differences between frequencies of observed phenotypes. *p <* 0.05 were considered significant; * *p <* 0.05, ** *p <* 0.01, *** *p <* 0.001, **** *p <* 0.0001.

Polo-mediated Cnb phosphorylation is important for Cnb’s timely mother to daughter centriole relocalization in mitosis (Gallaud *et al*., 2020). In human cells, Pp4 interacts with Centrobin (Réthi-Nagy *et al*., 2023) leading us to hypothesize that Pp4-mediated dephosphorylation of Centrobin could be required for mitotic Cnb relocalization. To test this model, we employed a phospho-mimetic fluorescent construct of Centrobin, where three of its Polo phosphorylation sites are mutated to glutamic acid, mimicking a phosphorylated state (w;;pUb-YFP::Cnb(T4E,T9E,S82E), (Januschke *et al*., 2013a)). Neuroblasts expressing this phospho-mimetic Centrobin mutant showed centriolar transfer defects similar to what we observed in *Pp4^Δ^* mutants (Figure 6C, E-G). Cnb was mostly localized to mother centrioles in prometaphase and metaphase but failed to shift efficiently to daughter centrioles in anaphase and telophase (Figure 6C). We found several instances with Cnb localized to both the mother and daughter centriole in meta-and anaphase neuroblasts (Figure 6C-H, Supplemental Figure 5B-E). In telophase, neuroblasts expressing phosphomimetic Centrobin displayed a similar variation of Cnb enrichment compared to *Pp4* mutants. 50% of Cnb phosphomimetic-expressing neuroblasts displayed Cnb^T4E,T9E,S82E^ enrichment on daughter centrioles, 40% displayed inverted Cnb^T4E,T9E,S82E^ enrichment on the mother centriole, and 10% showed symmetric Cnb^T4E,T9E,S82E^ enrichment (Figure 6F-H).

To confirm these results, we live-cell imaged *Pp4^Δ^* mutants expressing Cnb::EGFP and mCherry::Jupiter in late telophase and early anaphase. Since we found *Pp4^Δ^* mutant neuroblasts in telophase either devoid of Cnb or with Cnb on both centrioles with 3D-SIM imaging, we reasoned that these Cnb localization defects in mitosis should give rise to some self-renewed early interphase neuroblasts containing either two Centrobin-positive centrioles or two Centrobin-negative centrioles. In line with this hypothesis, we found 16% of *Pp4* mutant neuroblasts where Centrobin appeared on both centrosomes and 7% where Centrobin was undetectable on either centrosome. The remaining *Pp4^Δ^* mutant neuroblasts contained a normal single Cnb^+^ centriole (Figure 7A-F). Collectively, we conclude that Pp4 is required for efficient and timely transfer of Cnb from the mother to the daughter centriole in mitosis. Furthermore, in agreement with our earlier findings (Gallaud *et al*., 2020), the data suggests that Cnb’s timely relocalization from the mother to the daughter centriole in mitosis depends on Cnb’s phosphorylation status.

**FIGURE 7:**
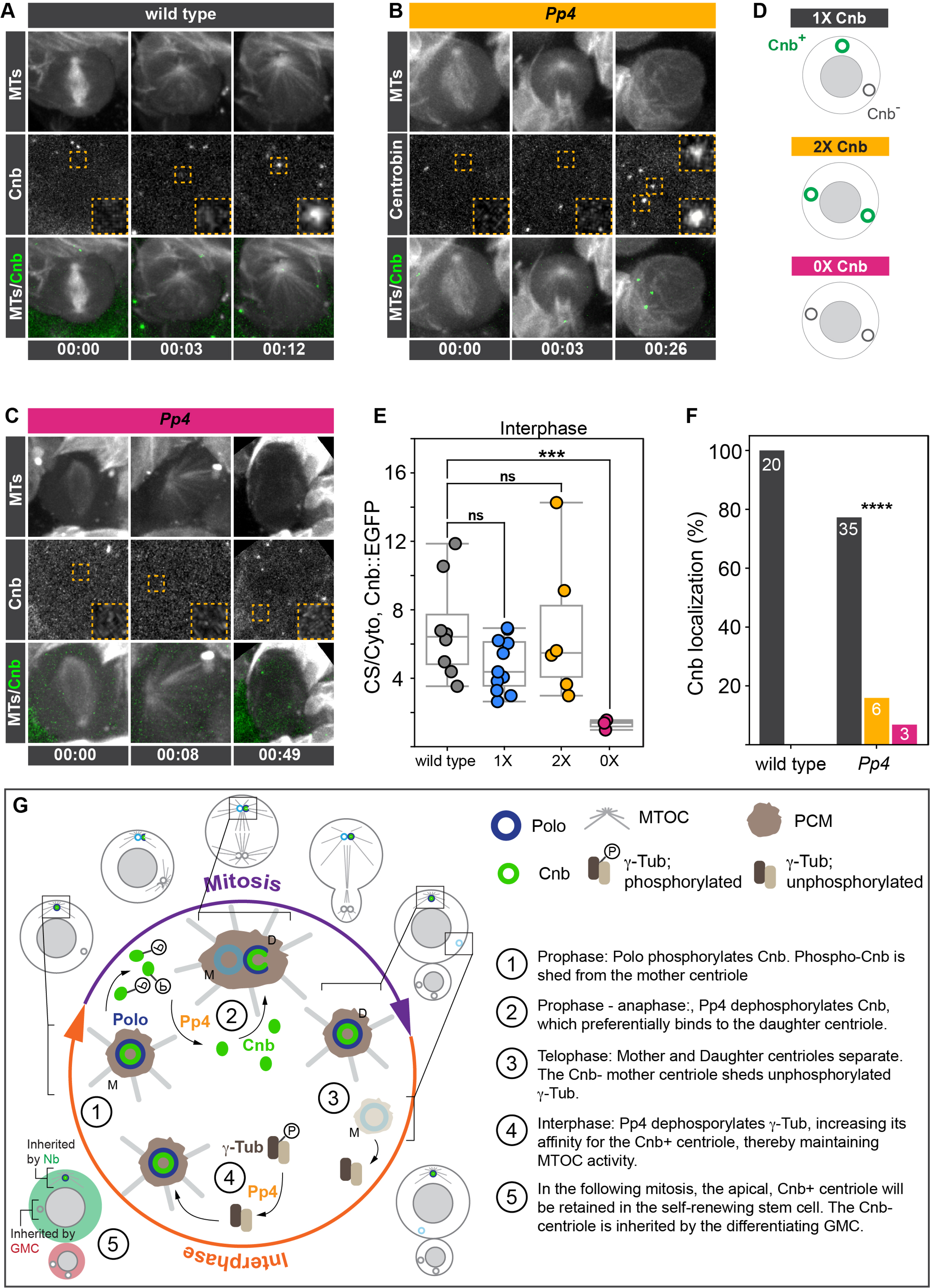
Pp4 is required for proper transfer of Centrobin. Representative image sequence of a **(A)** wild type or **(B, C)** two representative *Pp4^Δ^* mutant neuroblasts, expressing *worGal4, UAS-mCherry::Jupiter* (top row: white) and *Centrobin::GFP* (middle row: white; bottom row: green). Orange dashed boxes indicate apical centrosomes. High magnification images are shown in inserts on the bottom left in the middle row. **(B)** *Pp4^Δ^* mutant neuroblasts either contain Cnb on both centrosomes in early interphase or **(C)** is not detectable. **(D)** Schematic representation of the 0x, 1x and 2x Cnb phenotype. **(E)** Centrosome/Cytoplasm ratio of Centrobin intensity in wild type (gray), *Pp4^Δ^* with 1X Cnb (blue), *Pp4^Δ^* with 2X Cnb (yellow), and *Pp4^Δ^* with 0X Cnb (magenta). **(F)** Cnb localization frequencies are calculated based on (E) for wild type and *Pp4^Δ^* mutant neuroblasts. Numbers on bars denote number of measured neuroblasts. Cnb; Centrobin. MTs; microtubules. Scale bar denotes 5 µm. Statistical significance between normal distributions of data was determined via Student’s unpaired t-test to compare the differences in the distribution of collected data. Significance between non-parametric distributions of data was determined with a two-sided Mann-Whitney U statistical test. The chi-squared goodness-of-fit statistical test was done to test for differences between frequencies of observed phenotypes. *p <* 0.05 were considered significant; * *p <* 0.05, ** *p <* 0.01, *** *p <* 0.001, **** *p <* 0.0001. Time in: h:mins. **(G)** Proposed Model. Pp4 is needed to establish centriolar asymmetry in mitosis and to maintain MTOC activity on the Cnb^+^ centriole in interphase. In mitosis, Polo-mediated phosphorylation of Cnb removes it from the mother centriole but needs to be dephosphorylated by Pp4 for its deposition on the daughter centriole. In interphase, Pp4 dephosphorylates γTubulin, which incorporates into the Cnb^+^ centriole-containing centrosome. The Cnb^-^ centriole will shed PCM shortly after separating from the daughter centriole and loses MTOC activity. As a consequence of this centrosome asymmetry cycle, the Cnb^+^ centriole will be retained by the self-renewed neuroblast and the Cnb^-^ centriole will segregate into the differentiating GMC.

## DISCUSSION

Biased segregation of molecularly distinct centrosomes has been proposed to be important for asymmetric cell division and cell fate decisions (Sunchu and Cabernard, 2020b). However, the mechanisms underlying centrosome asymmetry remain unclear. Here, we show that in fly neural stem cells, the Protein Phosphatase 4 complex plays a dual role in centrosome asymmetry: in interphase, Pp4 is necessary to maintain MTOC activity on the centrosome destined to be inherited by the self-renewing neural stem cell. We also find that Pp4 is required for the timely localization of Cnb from the mother to the daughter centriole during mitosis.

Previous work revealed that fly neural stem cells contain two molecularly asymmetric interphase centrosomes: a younger, Cnb^+^ centriole and an older, Cnb^-^ centriole. The Cnb^+^ centriole retains the ability to nucleate microtubules, thereby tethering this centrosome to the apical neuroblast cortex. The Cnb^-^ centriole-containing centrosome downregulates PCM components such as centrosomin (Cnn) or γTubulin in interphase. The Cnb^-^ centrosome migrates through the cytoplasm in interphase and is destined to be inherited by the differentiating GMC in the next division. Thus, biased MTOC activity relies on the establishment of two molecularly distinct centrioles and is necessary for the asymmetric centrosome segregation.

We recently showed that Cnb asymmetry is established in mitosis, when neural stem cell centrioles duplicate. In prophase, Cnb starts to change its localization from the older mother centriole to the newly forming daughter centriole. The mechanisms are not entirely clear but include a downregulation of Cnb from the mother in prophase, a relocalization of the remaining Cnb from the mother to the daughter centriole between prometaphase and anaphase, and finally recruitment of new Cnb to the maturing daughter centriole in telophase and early interphase. The establishment of asymmetric Cnb localization in mitosis depends on the mitotic kinase Polo and Cnb’s phosphorylation status (Gallaud *et al*., 2020). Here we show that in addition to Polo, Pp4 is also required for the timely relocalization of Cnb from the mother to the daughter centriole in mitosis. In mitotic *Pp4^Δ^* mutants, we found neuroblasts showing either Cnb enrichment on the mother centriole, symmetrical distribution or no Cnb enrichment on either centriole. Pp4 and Centrobin have been shown to interact *in vitro* (Lipinszki *et al*., 2015a), suggesting that Cnb’s phosphorylation status is important for the timely relocalization in mitosis. Indeed, *polo* mutants, or unphosphorylatable Cnb show defects in Cnb relocalization to the daughter centriole in mitosis (Gallaud *et al*., 2020). In support of this finding, we demonstrate that phospho-mimetic Centrobin displayed transfer defects like those observed in *Pp4^Δ^* mutants. Accurate relocalization of Centrobin in mitosis ensures the formation of a Cnb^+^ centriole and one Cnb^-^ centriole, separating from each other in early interphase. Therefore, Cnb relocalization defects could either give rise to (1) two Cnb^-^, (2) two Cnb^+^ or (3) one Cnb^-^ and one Cnb^+^ centriole, with Cnb remaining on the older mother centriole. In support of our OMX data, and consistent with this model, our live cell imaging data showed *Pp4^Δ^* mutant neuroblasts with either two Cnb^+^ centrosomes or two Cnb^-^ centrosomes in interphase. The majority of live imaged *Pp4^Δ^* mutant neuroblasts showed asymmetric Cnb distribution, but we cannot resolve whether Cnb remained on the mother centriole or correctly localized to the daughter centriole. Based on these data, we propose that in mitosis, Polo is responsible for the removal of Centrobin from the mother centriole in prophase and prometaphase when Polo is strongly enriched on the mother centriole. We further propose that Pp4 is then required to deposit Centrobin onto the nascent daughter centriole, which may have higher affinity for unphosphorylated Cnb.

Pp4 is composed of a catalytic subunit Pp4c, as well as the two regulatory subunits Pp4r2, and Pp4r3, which provide substrate and localization specificity (Park and Lee, 2020). Loss of Pp4c, Pp4r2 or Pp4r3 all cause loss of MTOC activity in interphase, a phenotype resembling the loss of Cnb. Building a robust MTOC requires correctly organized PCM proteins, including *γ*Tubulin. *Drosophila* contains two *γ*Tubulin isoforms (*γ*Tubulin23C and *γ*Tubulin37C), which physically interact with each other (Wiese, 2008). We found that Pp4 is required for *γ*Tubulin23C and *γ*Tubulin37C localization at the apical centrosome and both isoforms are necessary for correct MTOC asymmetry. Previous studies revealed that *γ*Tubulin23C is required for all centrosomal function whereas *γ*Tubulin37C is required for meiotic spindle regulation and early oogenesis (Vázquez *et al*., 2008; Wiese, 2008). Our results suggest that the two isoforms are partially redundant with each other, since *γ*Tubulin single, but not double mutants still localize *γ*Tubulin to centrosomes. *In vitro* data further suggest that *γ*Tubulin is dephosphorylated by Pp4 and that constitutively phosphorylated *γ*Tubulin at Ser131 lowers microtubule nucleating activity (Alvarado-Kristensson *et al*., 2009; Voss *et al*., 2013). Ser131 is conserved in both Tubulin23C and *γ*Tubulin37C but mutating Serine 131 to non phosphorylatable Alanine in *γ*Tubulin37C is sufficient to maintain two active interphase MTOCs, suggesting that the dephosphorylated state is necessary for MTOC activity in fly neural stem cells. Cdk1 has also been implicated in *γ*Tubulin regulation, either via inhibiting Pp4 or by phosphorylating *γ*Tubulin. In both cases, preventing *γ*Tubulin dephosphorylation diminishes its microtubule nucleating activity (Alvarado-Kristensson *et al*., 2009). Consistent with this model, we found that loss of Cdk1, or Cdk1 mislocalization compromises interphase MTOC activity. Despite these results, we have no evidence to suggest that Pp4 dephosphorylates or CDK1 phosphorylates *γ*Tubulin’s Ser131 residue.

In conclusion, we propose that Pp4 dephosphorylates Cnb in early mitosis to aid its timely and correct relocalization from the mother to the nascent daughter centriole. Although we have not mapped the potential target sites, previously studied Polo-phosphorylation sites are necessary for the correct relocalization of Cnb. This could mean that Polo and Pp4 both regulate the same site in a sequential manner to first remove – via Polo mediated phosphorylation – and then add back Cnb – via Pp4 dephosphorylation (Figure 7G). Consistent with this model is the finding that in vertebrate cells, Polo mediates the removal of Centrobin from the mother centriole at the prometaphase – metaphase transition (Roux-Bourdieu *et al*., 2022). We further propose that Cdk1 and Pp4 regulate *γ*Tubulin in interphase to maintain robust MTOC activity on only the Cnb^+^ centriole. Although we have not directly implicated specific residues, the conserved Serine 131 residue seems to be important as its unphosphorylated state gives rise to two active MTOCs in interphase. The identification of the target residues and the spatiotemporal activity of Polo, Cdk1 and Pp4 will remain future challenges to provide mechanistic insight into centrosome asymmetry.

## MATERIALS AND METHODS

### Fly Strains

Mutant alleles, transgenes and fluorescent markers:

Pp4^Δ^/FM7C ActGFP (this work), worGal4, UAS-Cherry::Jupiter (Cabernard and Doe, 2009b), Asl::GFP/CyO (Blachon *et al*., 2008b), w^-^; 5X-UAS-Pp4::GFP/CyO, w^-^; 10X-UAS-Pp4[D85N, H115N]::GFP/CyO (Lipinszki *et al*., 2015b), y1,v1; P{y[+t7.7] v[+t1.8]=TRiP.JF02065}attP2 (Pp4r2 RNAi; BDSC26296), w^-^; P{ry[+t7.2]=neoFRT}82B flfl[795]/TM6B, Tb[+] (Sousa-Nunes *et al*., 2009), w^-^; P{w[+mC]=XP-U}Exel6170/TM6B, Tb[1] (Deficiency, removing Falafel; BDSC7649), ncd-γTubulin37C::GFP/TM6B,Tb (Hallen *et al*., 2008b, 2008a), γTubulin23C::GFP/TM6B,Tb (Mukherjee *et al*., 2020), w-; P{w[+mC]=XP-U}Exel6043/CyO (Deficiency, removing γTubulin37C; BDSC7525), w-; Df(2L)JS31, dpp[d-ho]/CyO (Deficiency, removing γTubulin23C; BDSC64238), w-; γTubulin23C[A6-2]/CyO, w-; γTubulin23C[A14-9]/CyO, w-; γTubulin23C[A15-2]/CyO (Vázquez *et al*., 2008), w-; γTubulin37C[3]/CyO (Wilson and Borisy, 1998b), w-; PBac{w[+mC]=RB}gammaTub37C[e00793]/CyO (Thibault *et al*., 2004), w-; γTubulin37C[S131A]/CyO Actin-GFP; MKRS/TM6B,Tb (this work), w-; γTubulin23C[A15-2], γTubulin37C[3]/CyO Actin-GFP; (this work), y1,v1; P{y[+t7.7] v[+t1.8]=TRiP.JF03004}attP2; (RNAi against Cdk1; BDSC28368), w-; IF/CyO-Actin-GFP; Cdk1::EGFP/TM6B, Tb (this work), Sas4::GFP/TM6B,Tb (Peel *et al*., 2007), w-; UAS-vhh4::GFP/TM6B,Tb (Caussinus *et al*., 2012), Cnb::GFP/TM6B,Tb (Gallaud *et al*., 2020), Polo::GFP/TM6B,Tb (Gallaud *et al*., 2020), w-; Asterless::Cherry/CyO (Conduit and Raff, 2010), w;;pUb-YFP::Cnb[T4E,T9E,S82E] (Januschke *et al*., 2013a), w; If/CyO; cnb e00267/TM6B, Tb (Deficiency removing Cnb; (Januschke *et al*., 2013b)). Experimental crosses were performed as outlined in Supplemental Table 2. All strains were raised on standard medium at 25°C, under a 12L:12D light cycle.

### Immunohistochemistry

The following antibodies were used for this study: Mouse anti-alpha-tubulin (Sigma, 1:2000, catalog number T6199), Guinea Pig anti-Asterless (Gifted from J. Raff, 1:20,000), mouse anti-gamma-tubulin (Millipore Sigma, 1:2000, catalog number SAB4701044). Secondary antibodies were from Molecular Probes and all done at 1:1000.

72 to 96 hours after egg laying, larval brains were dissected in Schneider’s insect medium (Sigma, catalog number S0146-100ML) for no more than 20 minutes. Briefly, dissection was done by removing the posterior end of the larvae and inverting the entire larvae. After inversion, the fat bodies and gastrointestinal tract were removed, and the axonal connections between the brain lobes and cuticle were severed. This resulted in an inverted cuticle devoid of all other organs save for the brain, which is connected to the cuticle via axonal connections with the ventral nerve cord. After dissection, samples were fixed in 4% paraformaldehyde in Schneider’s medium for 20 minutes on a rotator at room temperature. After fixing, the samples were washed at least three times with PBSBT (1X PBS, 0.1% volume/volume of Triton-X-100, and 1% weight/volume BSA) for one hour. Primary antibody dilution was prepared in 1X PBSBT, and samples were incubated in primary antibody solution for at least two days at 4°C with agitation. After primary labeling, samples were washed with PBSBT at least three times for 20 minutes each. Secondary antibody solution was prepared in 1X PBSBT, and samples were covered and again incubated for two days at 4°C with agitation. After incubation, the samples were washed three times with PBST (1X PBS, 0.1% volume/volume of Triton-X-100). During this final wash, mounting slides were prepared by affixing two glass coverslips onto a glass slide with nail polish to form a thin channel to hold the sample. Final dissections to remove the brains from the cuticles were done in PBSBT, and the brains were then transferred via pipette to equilibrate in Vectashield H-1000 mounting media (Vector laboratories). After equilibration, samples were transferred to the previously prepared slides and sealed with a glass coverslip and VALP (1:1:1 by weight mixture of Vaseline, Lanolin, and Parafin). Samples that were not imaged were stored in Vectashield at 4°C for up to several weeks.

### Live cell imaging

72 to 96 hours after egg laying, larval brains were dissected using microdissection scissors (Fine Science Tools catalog number 15003-08) (Segura and Cabernard, 2023) and forceps (Dumont #5, Electron Microscopy Sciences, item number 0103-5-PO) in Schneider’s medium supplemented with 10% bovine growth serum (BGS) (HyClone, item number SH30541.03) and transferred to chambered slides (Ibidi, catalog number 80826) for imaging. Live samples were imaged either with an Intelligent Imaging Innovations (3i) spinning disc confocal system, consisting of a Yokogawa CSU-W1 spinning disc unit and two Prime 95B Scientific CMOS cameras, on on an Andor BC43 benchtop confocal microscope. A 60x/1.4NA oil immersion objective mounted on a both microscopes was used for imaging. Live imaging voxels are 0.22 × 0.22 × 1 μm (60x/1.4NA spinning disc). Temporal resolution varied between 30 seconds and three minutes per frame.

### 3D structured illumination microscopy

3D-SIM was performed on fixed brain samples using a DeltaVision OMX-Blaze system (version 4; GE Healthcare), equipped with 405-, 445-, 488-, 514-, 568-, and 642-nm solid-state lasers. Images were acquired using a Plan Apo N 60x, 1.42 NA oil immersion objective lens (Olympus) and 4 liquid-cooled sCMOs cameras (pco.edge 5.5, full frame 2560 × 2160; PCO). Exciting light was directed through a movable optical grating to generate a fine-striped interference pattern on the sample plane. The pattern was shifted laterally through 5 phases and 3 angular rotations of 60° for each z section. Optical z sections were separated by 0.125 μm. The laser lines 405, 488, 568, and 642 nm were used for 3D-SIM acquisition. Exposure times were typically between 10 and 120 ms, and the power of each laser was adjusted to achieve optimal intensities of between 5,000 and 8,000 counts in a raw image of 15-bit dynamic range at the lowest laser power possible to minimize photobleaching. Multichannel imaging was achieved through sequential acquisition of wavelengths by separate cameras.

### 3D-SIM image reconstruction

Raw 3D-SIM images were processed and reconstructed using the DeltaVision OMX SoftWoRx software package (version 6.1.3, GE Healthcare; Gustafsson, M. G. L. 2000). The resulting size of the reconstructed images was of 512 × 512 pixels from an initial set of 256 × 256 raw images. The channels were aligned in the image plane and around the optical axis using predetermined shifts as measured using a target lens and the SoftWoRx alignment tool. The channels were then carefully aligned using alignment parameter from control measurements with 0.5-μm diameter multispectral fluorescent beads (Invitrogen, Thermo Fisher Scientific).

### Analysis

All analyses were done in Imaris 10.1 and ImageJ 1.54. Imaris was used to determine the mean and integrated fluorescent density present at centrosomes. ImageJ was used to generate line scans of centrosomal intensity. The Imaris spot tool was used to measure the mean and integrated fluorescent density present at centrosomes in interphase and mitosis. Interphase was defined as at least 30 minutes before nuclear envelope breakdown (NEB), and mitosis was defined as five to seven minutes before NEB, which was defined as the first frame where microtubule penetration into the nucleus was observed. A 1.83x1.83x1.83 μm (xyz) spot was centered on the centrosome using the location of centriolar signal from either Asterless-GFP or Centrobin-GFP, and then used to measure either the total sum or average fluorescence intensity of either the GFP (Asterless, Centrobin, or Polo) or RFP (Cherry::Jupiter, microtubules) channels. For *γ*Tubulin analysis, spot sizes were adjusted to either 1x1x2 μm (xyz) for the BC43 andor microscope or 1.5x1.5x2 μm for the 3i Nikon microscope. To normalize against cytoplasmic intensity, one spot of the same size was placed in the cytoplasm. The average fluorescence intensity of both centrosomal and cytoplasmic measurements were then used to normalize centrosomal signal to cytoplasmic signal, such that:

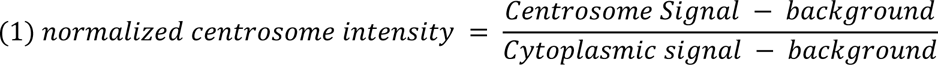

Where “background” is defined as the minimum value assigned to pixels by the microscope camera. Similarly, asymmetry indices were generated by normalizing the apical centrosome to the basal centrosome for markers of interest, such that:

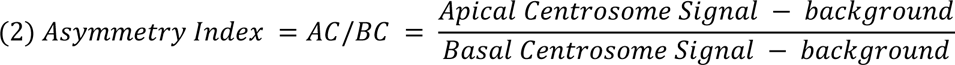

Where AC and BC refer to the apical and basal centrosomes respectively. These imaris spots were also used to infer centrosome-to-centrosome positioning based on their X, Y, and Z coordinates. The distance between centrosomes was calculated based on the equation for the distance between two points in 3D space, such that if centrosome 1 is defined as (x_1_, y_1_, z_1_) and centrosome 2 is defined as (x_2_, y_2_, z_2_):

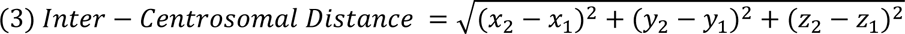

This calculation was also applied if the distance between two points was measured, such as the distance between the apical centrosome and the apical cortex.

Imaris Measurement Points were used to determine the angle of movement between the position of a centrosome in interphase and its ultimate location at metaphase. This was done by placing point A on the centrosome in interphase, point B on the center of the cell, and point C on the centrosome in metaphase, and the measurement of the angle formed by ∠ABC was recorded and plotted.

ImageJ was used to generate intensity plots of centrosomal signal. Using the same cell cycle timepoints as above, the center-most slice of the centrosome was identified. Following this, a summed projection of the center-most slice plus the slices directly above and below the center was generated. Using this summed projection, a line of at least five microns was drawn through the center point of the centrosome (again, based on centriolar markers such as Asterless and Centrobin). This line was then used to generate a line plot of intensity, which was recorded and saved as a csv file for analysis.

### Centriolar Age Measurement

To determine centriolar age, Asl intensity was used as a reference. The contours of nonoverlapping centrioles were drawn in ImageJ based on Asl signal and saved as a region of interest (ROI) in imageJ. Total Asl intensity was then used to determine centriolar age because daughter centrioles have lower intensity than mother centrioles (Gallaud *et al*., 2020). The same ROIs were then used to measure total pixel intensity for markers of interest (e.g., γTubulin37C::GFP, Cnb::EGFP). Asymmetry ratios for markers of interest were then determined by dividing total daughter centriole intensity with total mother centriole intensity.

### Definition of Daughter versus Mother enrichment of Centrobin

To determine the enrichment of Centrobin on centrioles, we used the raw integrated density of Centrobin signal observed in the cytoplasm (B_signal_), on the daughter centriole (D_signal_), and on the mother centriole (M_signal_). To determine the variation of signal in the background, we measured 60 regions of interest in the cytoplasm of one WT cell and used the resulting values to determine the percent standard deviation observed. The following logic gates were then used in Python to determine enrichment:

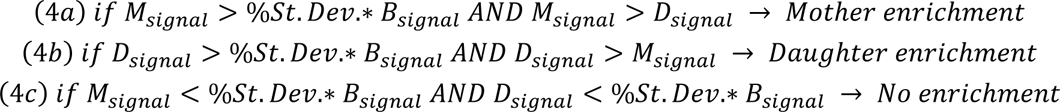

The resulting determinations of enrichment were then used to bin the data for visualization.

### Determination of Daughter:Mother Centriole Ratios

To determine the ratio of Daughter to Mother centriole signal of either Centrobin or Asterless, the Integrated Density of signal was used in the following equation:

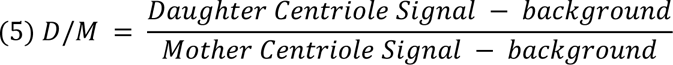

In instances where the background signal exceeded that of either the mother or daughter centriole (Supplemental Figure 6A-6E), the following was used instead:

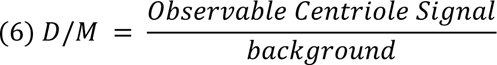

Where *Observable Centriole Signal* is the signal of either the Daughter or the Mother.

### Generation of the PP4 19C Null Allele

CRISPR target sequences for the PP4-19c locus were identified using the FlyCRISPR Target Finder tool (https://flycrispr.org/target-finder/). Two guide RNAs were identified that flank the PP4-19c gene to generate a precise deletion without affecting neighboring genes: 5’ TTCGCTTCGCTTTTGCAGGC ’3 and 5’ ACTTGTTGAGATAGTCCATG ’3. gRNAs were cloned into the pBbsI-U6-gRNA expression vector (p-U6-BbsI-chiRNA; cat# 45946, Addgene) using BbsI mediated cloning following a standard protocol (https://flycrispr.org/protocols/).

Primer: 5’<u>CTTC</u>**G**TTCGCTTCGCTTTTGCAGGC’3, 5’<u>CTTC</u>**G**ACTTGTTGAGATAGTCCATG’3

A donor plasmid (pHD-DsRed-attP, cat# 51019, Addgene) was used for homology-directed repair (HDR) to replace the PP4-19c locus with a marker cassette. Approximately 1 kb homology sequences upstream of the 5′ CRISPR site and downstream of the 3′ site were amplified by PCR from genomic DNA and cloned into the donor vector carrying a 3xP3>dsRed cassette to enable identification of successful replacement events via fluorescent screening. The donor vector was assembled using standard molecular cloning techniques and verified by sequencing.

Primer 5’Homology arm (<u>AarI</u>): 5’tgca<u>CACCTGC</u>gatc<u>ctac</u>ATGAGGCTATTTCGCAATG’3, 5’attc<u>CACCTGC</u>tgaa<u>tcgc</u>CCTTGTCTCTGTGATCTC’3

Primer 3’Homology arm (<u>SapI</u>): 5’ cgag<u>GCTCTTC</u>a<u>gac</u>TGCAGAGAAGGAAGACGC’3, 5’ tagt<u>GCTCTTC</u>a<u>tat</u>TGCAAAAGCGAAGCGAAC’3

gRNA vectors and the donor plasmid were co-injected into vasa-Cas9 embryos (BL#51324).

Injected embryos were raised to adulthood and crossed to a balancer stock. F1 progenies were screened for dsRed fluorescence in the eye using a fluorescence stereomicroscope. Fluorescent-positive individuals were selected as potential PP4-19c deletion mutants.

### Molecular Verification of PP4 19C null

Genomic DNA was extracted from fluorescent-positive heterozygous F1 individuals. PCR was performed using primers flanking the PP4 19C locus to confirm successful replacement of the target region with the 3xP3>dsRed cassette (expected size 1.9 kb for control and 1.7 kb and 1.9 kb for heterozygous mutants. PCR products were validated by Sanger sequencing.

Primers: 5’CAGGCGAACTCAAAAACCG’3, 5’GGTAGAGGTAAACAGATGG’3

### Generation of PP4 19C transgenic flies

The full length PP4-19c ORF was amplified from the PP4-19c cDNA clone) RE58406 obtained from the Drosophila Genomic Research Center (Indiana, USA) using the following primers: 5’<u>CACC</u>ATGTCCGACTACAGCGAC’3 and 5’TTAGAGAAAGTAGTCCGCCTG’3

The ORF was cloned into pENTR using TOPO cloning (Invitrogen) and exchanged into the pUASattB-10xUAS and 5xUAS destination vectors with either no tag, a C-terminal EGFP or an N-terminal EGFP tag. Phosphatase dead versions of PP4 19C were generated using entry vectors carrying PP4 19C^D85N,H115N^ and PP4 19C^D85N^, kindly provided by Dr. Marcin R. Przewloka (Lipinszki *et al*., 2015a). All constructs were verified by sequencing and injected into the attP40 genomic landing site (BestGene Inc, California, USA).

### Generation of *γ*Tubulin phospho-mutants

The *γ*γTubulin37C coding sequence region was amplified using the following primers:

5’CTACATTCCTCGTGGG3’

5’GCCGTGCTTTAGCGTA3’

Serine 131 of gamma tubulin37C was mutated to Alanine via CRISPR-mediated homology-directed repair. We introduced a double-stranded break via the CRISPR target CGGCTTTAGCCAGGGTGAGAAGG, 50 base pairs (bp) away from Serine131. We then designed rescue templates that introduced a mutated Serine131 (changing TCG to GCG). Rescue templates carrying the specific site mutation could not be generated via PCR and were instead synthesized by GeneWiz. Lowercase bold bases denote homology with the dsRed backbone:

5’**taaagatctccatgcataaggcgcgcc**TCCCCTTTGCAGTTGTACAACCAGGAGAATGTGTTCCTCTCCAAGCACGGCGGTGGAGCAGGAAACAACTGGGCGTCCGGCTTTAGCCAGGGTGAGAAG**a**TGCAGGAGGAGGTGTTCGACATTCTGGACCGCGAGGCCGATGGCAGCGAT**g*CG***CTGGAGGGATTCGTGC TCTG 3’ (homology with AscI, underline bold denotes mutated PAM site, bold underlined italic denotes mutated Serine 131)

The resulting rescue homology templates were then inserted into a EcoRI/NdeI/XhoI/AscI digested dsRed reporter plasmid via an infusion reaction kit. The following primers were used to generate homology arms for insertion into the dsRed vector:

5’**tcgctgaagcaggtggaatt**CTTCTCTGCGATTTTTACG3’ (upstream homology, EcoRI)

5’**tgatcgcaggtgtgcatatg**AGTTTTTTAAATGAAATACATAAAA 3’ (upstream homology, Nde1)

5’CTGGAGGGATTCGTGCTCTGCCACTCGATAGCCGGCGGAA3’ (downstream homology, homology with GeneWiz synthesized fragment)

5’ **attgacggaagagcctcgag**GCCAAGGCCCTCTTGAAGAG 3’ (downstream homology, XhoI)

Homology rescue templates were designed such that the dsRed reporter would be inserted into an intron upstream of Serine131 as to not interfere with viability. The plasmid and Cas9 delivery system were then injected into flies by BestGene, and dsRed was used to select for positive transformants. Once isolated and balanced over balancer chromosomes CyO-Actin::GFP, the dsRed cassette was floxed out of the newly generated line by crossing against a Cre-expressing line. Final transformants were verified with PCR and sequencing with primer 5’ AGCACCACCACACTGT3’ to confirm the presence of the desired mutation in the final line.

### Generation of Cdk1::EGFP Crispr line

Target specific sequences with high efficiency were chosen using the CRISPR Optimal Target Finder (http://tools.flycrispr.molbio.wisc.edu/targetFinder/), the DRSC CRISPR finder (http://www.flyrnai.org/crispr/), and the Efficiency Predictor (http://www.flyrnai.org/evaluateCrispr/) web tools. The gRNA was cloned into the pBbsI-U6-gRNA expression vector (p-U6-BbsI-chiRNA; cat# 45946, Addgene) using BbsI mediated cloning following a standard protocol (https://flycrispr.org/protocols/). The following primers were used: Cdk1_C_sgRNA_sense: CTTCGATGCTCCAAAATGTCCTTGG; Cdk1_C_sgRNA_antisense: AAACCCAAGGACATTTTGGAGCATC. 1 kb homology arms flanking the insertion site were cloned into the previously described pHD-DsRed-attP-EGFP::SspB vector using the following primers: Cdk1_C_LHA_203_FWD: CTGGGCCTTTCGCCCGGGCTGAAGATTTTGCAATATTGTTTTT; Cdk1_C_LHA_203_REV: CTCCAAAATGTCCTTGGCtGAAATGCGATGAACTGGATCGT.

### Definition of Statistical Tests and Replicates

Data was collected from at least two independent experiments. Statistical analyses and visualization were done in Python. Statistical significance between normal distributions of data was determined via Student’s unpaired t-test to compare the differences in the distribution of collected data. Significance between non-parametric distributions of data was determined with a two-sided Mann-Whitney U statistical test. The chi-squared goodness-of-fit statistical test was done to test for differences between frequencies of observed phenotypes. *p <* 0.05 were considered significant; * *p <* 0.05, ** *p <* 0.01, *** *p <* 0.001, **** *p <* 0.0001.

### Data Visualization

Data was visualized in Python 3.5 Jupyter notebooks. Visualizations were generated using Matplotlib and Seaborn. Statistical tests were done using Scipy. Code, raw data and data used for plotting can be found here: https://github.com/Roberto-Carlos-Segura/PP4_paper_2024.

## Acknowledgements

We thank Cayetano Gonzalez, Tomer Avidor-Reiss and Jordan Raff for fly stocks and antibodies, Jeff Rasmussen and members of the Cabernard laboratory for helpful discussions and comments. This work was supported by the National Institutes of Health (R35GM148160 to CC) and the Ford Foundation (to CRS). Stocks were obtained from the Bloomington Drosophila Stock Center (BDSC), which is supported by grant P40OD018537 from the NIH Office of Research Infrastructure Programs (ORIP) in collaboration with the National Institute of General Medical Sciences (NIGMS), National Institute of Neurological Disorders and Stroke (NINDA) and National Institute of Child Health and Human Development (NICHD).

---

**Supplemental Figure 1: Pp4r2 and Falafel are required for MTOC asymmetry in interphase**

Representative image sequences of **(A)** wild type, **(B)** *flfl[795]*, **(E, F)** *Pp4r2 RNAi* expressing neuroblasts. All neuroblasts also express *worGal4, UAS-mCherry::Jupiter* (top row and middle row: white) and *Asterless::GFP* (bottom row: green). Microtubule intensity at the **(C)** apical centrosome normalized to cytoplasmic signal or **(D)** apical to basal centrosome intensity ratios in wild type (gray) and *flfl[795]* mutants (blue). Microtubule intensity at the **(G)** apical centrosome normalized to cytoplasmic signal or **(H)** apical to basal centrosome intensity for wild type (gray) and *Pp4r2 RNAi* expressing neuroblasts (blue). Asl; Asterless. MTs; microtubules. Scale bar denotes 5 µm. Statistical significance between normal distributions of data was determined via Student’s unpaired t-test to compare the differences in the distribution of collected data. Significance between non-parametric distributions of data was determined with a two-sided Mann-Whitney U statistical test. *p <* 0.05 were considered significant; * *p <* 0.05, ** *p <* 0.01, *** *p <* 0.001, **** *p <* 0.0001. Time in: h:mins.

**Supplemental Figure 2: *γ*Tubulin mutants show a loss of MTOC asymmetry**

Representative images of **(A)** *γTub23C^[A6-2]^*, **(B)** *γTub23C^[A14-9]^*, **(C)**, *γTub23C^[A15-2]^*, and **(D)** *γTub37C^[e00793]^* mutants expressing *worGal4, UAS-mCherry::Jupiter* and *Sas4::GFP*. Orange and magenta arrrowheads and dashed boxes denote the apical (AC) and basal centrosome (BC), respectively. **(E)** Phenotypic penetrance in percent. wild type (WT); one active MTOC; 0x: no active MTOC; 2x: 2 active MTOCs. MTs; microtubules. Scale bar denotes 5 µm. Time in: h:mins.

**Supplemental Figure 3: γTub23C and γTub37C are redundant for MTOC formation in *Drosophila* neuroblasts**

Representative **(A)** interphase and **(B)** metaphase images of wild type, *γ*Tubulin37C^[3]^, *γ*Tubulin23C^[A14-9]^, and *γ*Tubulin37C^[S131A]^ mutants stained with anti-Asterless (Asl; magenta, top and bottom row) and *γ*Tubulin (green, middle and bottom row). **(C)** AC/Cytoplasm or **(D)** BC/Cytoplasm ratios for wild type (grey), *γ*Tubulin37C^[3]^ (blue), *γ*Tubulin23C^[A14-9]^ (magenta), and *γ*Tubulin37C^[S131A]^ (orange) mutants in interphase and metaphase. **(E)** Representative images of wild type and *γ*Tubulin23C^[A15-2]^, *γ*Tubulin37C^[3]^ double mutant stained with anti-Asl (magenta, top row) and *γ*Tubulin (green, middle and bottom row). High magnification images of *γ*Tubulin at the apical (yellow dashed box) and basal (blue dashed box) centrosome are shown in the bottom row. Left column shows representative images in interphase, and the right column shows representative images at metaphase. **(F)** Representative images of wild type and *γ*Tubulin23C^[A15-2]^, *γ*Tubulin37C^[3]^ recombinants with anti-αTubulin (gray, top row) and anti-Asl (magenta, second row). Left column shows representative images in interphase, and the right column shows representative images at metaphase. Asl; Asterless. MTs; microtubules. Scale bar denotes 5 µm. Statistical significance between normal distributions of data was determined via Student’s unpaired t-test to compare the differences in the distribution of collected data. Significance between non-parametric distributions of data was determined with a two-sided Mann-Whitney U statistical test. *p <* 0.05 were considered significant; * *p <* 0.05, ** *p <* 0.01, *** *p <* 0.001, **** *p <* 0.0001.

**Supplemental Figure 4: 3D-SIM image analysis**

**(A)** plots of individual values of Centrobin intensity at the daughter centriole (blue), mother centriole (yellow) and background (grey) for *Pp4^Δ^* mutant interphase neuroblasts. Background values (black lines) were used to evaluate Cnb intensity levels. If background intensity exceeded detectable Cnb on the mother or daughter centriole, the ratio was arbitrarily set to ‘1’ and were also counted towards the pool of neuroblasts with no Cnb signal on either centrosome. **(B-E)** representative scatter plots (top) and bar plot of frequency (bottom) of wild type (grey), *Pp4^Δ^* mutant (blue), and YFP::Cnb^T4E,T9E,S82E^ expressing (orange) neuroblasts in prophase **(A)**, prometaphase **(B)**, metaphase **(C)**, anaphase **(D)**, and telophase **(E)**. Statistical significance between normal distributions of data was determined via Student’s unpaired t-test to compare the differences in the distribution of collected data. Significance between non-parametric distributions of data was determined with a two-sided Mann-Whitney U statistical test. The chi-squared goodness-of-fit statistical test was done to test for differences between frequencies of observed phenotypes. *p <* 0.05 were considered significant; * *p <* 0.05, ** *p <* 0.01, *** *p <* 0.001, **** *p <* 0.0001.

**Movie 01: wild type neuroblast expressing worGal4, UAS-mCherry::Jupiter and Asl::GFP**

Wild type control larval neuroblast expressing the microtubule marker mCherry::Jupiter (in white in left and merge) and the centriolar marker Asterless::GFP (green in merge). Orange and magenta arrowheads denote the apical centrosome (AC) and basal centrosome (BC), respectively. Note that the AC remains active and anchored in the apical region of the cell throughout interphase (0:24:00). The BC matures and forms an MTOC at the onset of mitosis (0:42:00), after which the cell forms a bipolar spindle and divides. This division gives rise to one self-renewed neuroblast, and one differentiating GMC (0:54:00). Time scale is h:mm:ss, and the scale bar is 3 μm.

**Movie 02: *Pp4* mutant neuroblast expressing worGal4, UAS-mCherry::Jupiter and Asl::GFP**

*Pp4* mutant larval neuroblast expressing the microtubule marker mCherry::Jupiter (in white in left and merge) and the centriolar marker Asterless::GFP (green in merge). Orange and magenta arrowheads denote the apical (AC) and basal centrosome (BC), respectively. Note that both the AC and BC lack MTOC activity throughout interphase (0:15:00). Both centrosomes mature and form an MTOCs at the onset of mitosis (1:19:00), after which the cell forms a bipolar spindle and divides. Time scale is h:mm:ss, and the scale bar is 3 μm.

**Movie 03: wild type neuroblast expressing worGal4, UAS-mCherry::Jupiter and *γ*Tubulin23C::GFP**

Wild type control larval neuroblast expressing the microtubule marker mCherry::Jupiter (in white in merge on the right) and *γ*Tubulin23C::GFP (white on the left; green in merge). Orange and magenta arrowheads denote the apical (AC) and basal centrosome (BC), respectively. Note that *γ*Tubulin23C::GFP localizes to the AC throughout interphase (0:23:59). The BC matures and recruits *γ*Tubulin23C::GFP at the onset of mitosis (0:39:59), after which the cell forms a bipolar spindle and divides. Time scale is h:mm:ss, and the scale bar is 3 μm.

**Movie 04: *Pp4* mutant neuroblast expressing worGal4, UAS-mCherry::Jupiter and *γ*Tubulin23C::GFP**

*Pp4* mutant larval neuroblast expressing the microtubule marker mCherry::Jupiter (in white in merge on the right) and *γ*Tubulin23C::GFP (white on the left; green in merge). Orange and magenta arrowheads denote the apical (AC) and basal centrosome (BC), respectively. Note that the AC has reduced *γ*Tubulin23C::GFP throughout interphase (0:04:00). Both centrosomes mature and recruit *γ*Tubulin23C::GFP at the onset of mitosis (0:36:00), after which the cell forms a bipolar spindle and divides. Time scale is h:mm:ss, and the scale bar is 3 μm.

**Movie 05: **γ*Tubulin37C^[S131A]^* mutant neuroblast expressing worGal4, UAS-mCherry::Jupiter and Sas4::GFP**

**γ*Tubulin37C^[S131A]^* mutant larval neuroblast expressing the microtubule marker mCherry::Jupiter and the centriolar marker Sas4::GFP. Orange and magenta arrowheads denote the apical (AC) and basal centrosome (BC), respectively. Note that after division, the AC and BC separate from one another but maintain distinct MTOCs (0:35:59). These MTOCs remain active throughout interphase and increase upon entry into mitosis. Time scale is h:mm:ss, and the scale bar is 3 μm.

